# Wnt signaling promotes inflammation and EMT-associated gene expression in mesenchymal TNBC

**DOI:** 10.1101/2025.07.04.663027

**Authors:** Ramón García-Areas, Elodie Girard, Hamza Lasla, Virginie Raynal, Amit Kumar Pandey, Nicolas Servant, Pascal Jézéquel, Thierry Dubois

## Abstract

Aberrant activation of the Wnt signaling pathway in triple-negative breast cancer (TNBC) is linked to treatment resistance and recurrence, yet its role in tumoral heterogeneity remains unclear. We developed Wnt reporter cell lines from two mesenchymal TNBC models (MDA-MB-231 and MDA-MB-436) using a Tcf/Lef-eGFP vector and observed intra- and inter-tumoral variation in Wnt activity. Paired Wnt-positive and Wnt-negative TNBC cell lines were established and profiled by RNA-Seq. Integrative analyses revealed that Wnt-positive cells consistently upregulate genes involved in epithelial-to-mesenchymal transition, inflammation (e.g., IL6/JAK/STAT3, TNFα via NF-κB), and extracellular matrix remodeling. A 55-gene Wnt signature common to both nutrient conditions captured these features. Wnt-related gene sets were also enriched in the Mesenchymal-like Immune-Altered subtype of 699 primary TNBC tumors. These findings highlight the role of basal Wnt activity in driving pro-tumorigenic transcriptional programs in TNBC and provide new insight into its contribution to subtype-specific disease features.

## Introduction

Triple-negative breast cancer (TNBC) is the most aggressive breast cancer compared to other breast cancer subgroups^1^. TNBC tumors exhibit a lack of expression of estrogen and progesterone receptors, as well as an absence of gene amplification of *HER2*^2^. It has been estimated that TNBC patients constitute about 15% of the overall breast cancer patient population^3^. Paradoxically, patients diagnosed with TNBC often have a higher initial response rate to chemotherapy treatments compared to patients with other types of breast cancer^4^. This initial enhanced response may be attributed to TNBC tumors often being highly proliferative, and hence, being more responsive to chemotherapies targeting rapidly dividing cells. TNBC tumors also exhibit a complex cellular composition that yields strong intra- and inter-tumoral heterogeneity^5^. Several TNBC subtypes have been identified^6–8^, and our analysis of RNA-sequencing (RNA-Seq) data obtained from 699 TNBC led to the classification of four main distinct TNBC subtypes: Luminal Androgen Receptor (LAR), Mesenchymal-like Immune-Altered (MLIA), Basal-like Immune-Activated (BLIA), and Basal-like Immune-Suppressed (BLIS)^9^. The tumoral heterogeneity in TNBC has been identified as a potential factor influencing treatment resistance, relapse, and the development of metastatic lesions, which contributes to poor clinical outcomes^10,11^.

The pronounced tumor heterogeneity and dynamic treatment responses characteristic of TNBC have posed significant challenges in the identification of overarching therapeutic targets for TNBC. Nonetheless, notable progress has been achieved with the use of poly(ADP-ribose) polymerase (PARP) inhibitors for the treatment of TNBC harboring mutations in the *BRCA1* or *BRCA2* genes^12,13^. Concurrently, the rapidly growing field of immunotherapy has become a compelling therapeutic approach for combating highly aggressive solid tumors such as TNBC^14^. The combination therapy involving the anti-PD-L1 antibody atezolizumab and nab-paclitaxel was previously granted approval for the treatment of locally advanced or metastatic TNBC^15^. While the approval of atezolizumab plus nab-paclitaxel for TNBC was voluntarily withdrawn in 2021, pembrolizumab remains an approved option, used with chemotherapy for PD-L1-positive metastatic TNBC and as neoadjuvant/adjuvant therapy for high-risk, early-stage TNBC based on KEYNOTE-355 and KEYNOTE-522 trials^16,17^. However, given the heterogeneous nature of TNBC, further research into the mechanisms regulating immune responses is critical for the development of more effective immunotherapeutic strategies. Given the limited treatment options against TNBC, there is significant interest in deciphering the underlying signaling pathways that sustain TNBC tumorigenesis^18^, such as the Wnt signaling pathway, which was originally discovered as a mediator of mammary tumorigenesis^19^. This pathway plays a crucial role in promoting tumorigenesis and metastasis in TNBC^20^, and notably, it has been shown to mediate cancer cell stemness^21^ and resistance to chemotherapies and immunotherapies^22^. Because targeting the Wnt pathway could overcome treatment resistance and relapse, our group is actively evaluating it as an actionable target in TNBC^23–26^.

Activation of the β-catenin-dependent (canonical) Wnt pathway occurs through the binding of Wnt ligands to Wnt transmembrane receptors, for example, Frizzled receptors (e.g. FZD7) and co-receptors low-density-lipoprotein-related proteins 5 and 6 (LRP5/6)^27^. In various cancer types, including melanoma, colorectal, and pancreatic cancer, aberrant Wnt activity is often driven by mutations in Wnt pathway components^28^. In contrast, aberrant Wnt signaling in TNBC has been associated with an overexpression of Wnt ligands and Wnt receptors such as LRP5, LRP6, and FZD7^24,29^. Upon ligand-receptor engagement, the canonical Wnt pathway inhibits β-catenin degradation by the destruction complex, leading to its cytosolic accumulation and nuclear translocation^27^. In the nucleus, β-catenin associates with T-cell factor/lymphoid enhancer factor (Tcf/Lef) transcription factors, including TCF7, TCF7L1, TCF7L2, and LEF1, to co-activate the expression of Wnt target genes, thereby driving the Wnt transcriptional program^30^. The identification of a human Tcf/Lef binding DNA sequence (5′-AGATCAAAGG-3′)^31^, known as a Wnt Response Element (WRE), enabled the development of Tcf/Lef reporter vectors containing multiple WRE sequences linked to a reporter gene (e.g., Firefly luciferase or eGFP)^32,33^. These Tcf/Lef reporter assays have provided robust readouts to detect Wnt activity, although β-catenin-independent mechanisms may also contribute to Wnt signaling^34,35^.

Given the importance of Wnt activity in TNBC and the need to model tumoral heterogeneity, we developed a system to measure Wnt activity in TNBC cells while accounting for both intra- and inter-tumoral heterogeneity. To achieve this, we established Wnt activity reporter TNBC cell lines that utilize eGFP as a readout for Wnt activity. Specifically, we stably transfected two TNBC cell lines, MDA-MB-231 and MDA-MB-436, with the 7TGP Tcf/Lef vector (7xTcf-eGFP). Subsequently, two distinct populations, Wnt-positive and Wnt-negative, were isolated based on eGFP expression using fluorescence-activated cell sorting (FACS). These cell lines were cultured under low (0.5%) or standard (10%) serum conditions, and their transcriptomic profiles were determined by RNA sequencing (RNA-Seq). Bioinformatic analyses identified genes and signaling pathways associated with Wnt activity in these TNBC cells, and their expression levels were subsequently evaluated in a large cohort of TNBC patients.

## Results

### Wnt reporter activity reveals inter- and intra-tumoral heterogeneity in TNBC cell lines

The Wnt pathway has been shown to be activated in TNBC and to regulate tumorigenic processes in this highly aggressive type of breast cancer. To characterize Wnt activity while accounting for TNBC heterogeneity, we utilized the mesenchymal MDA-MB-231 and MDA-MB-436 TNBC cell lines, exhibiting enriched expression of WNT-related genes^6^. To assess Wnt activity, we used the 7TGP Tcf/Lef Wnt activity reporter vector^33^, which contains seven repeats of a consensus Tcf sequence (5′-AGATCAAAGG- 3) upstream of an eGFP reporter sequence, and a puromycin N-acetyltransferase selection cassette (Fig. 1a). The Tcf sequence in the 7TGP vector is considered as WRE and shares high homology with the consensus sequences bound by LEF1, TCFL1, and TCFL2 (Fig. 1b). Stable MDA-MB-231^7TGP^ and MDA-MB-436^7TGP^ cell lines were obtained by transfecting the 7TGP vector followed by puromycin selection (Fig. 1c).

**Figure 1.**
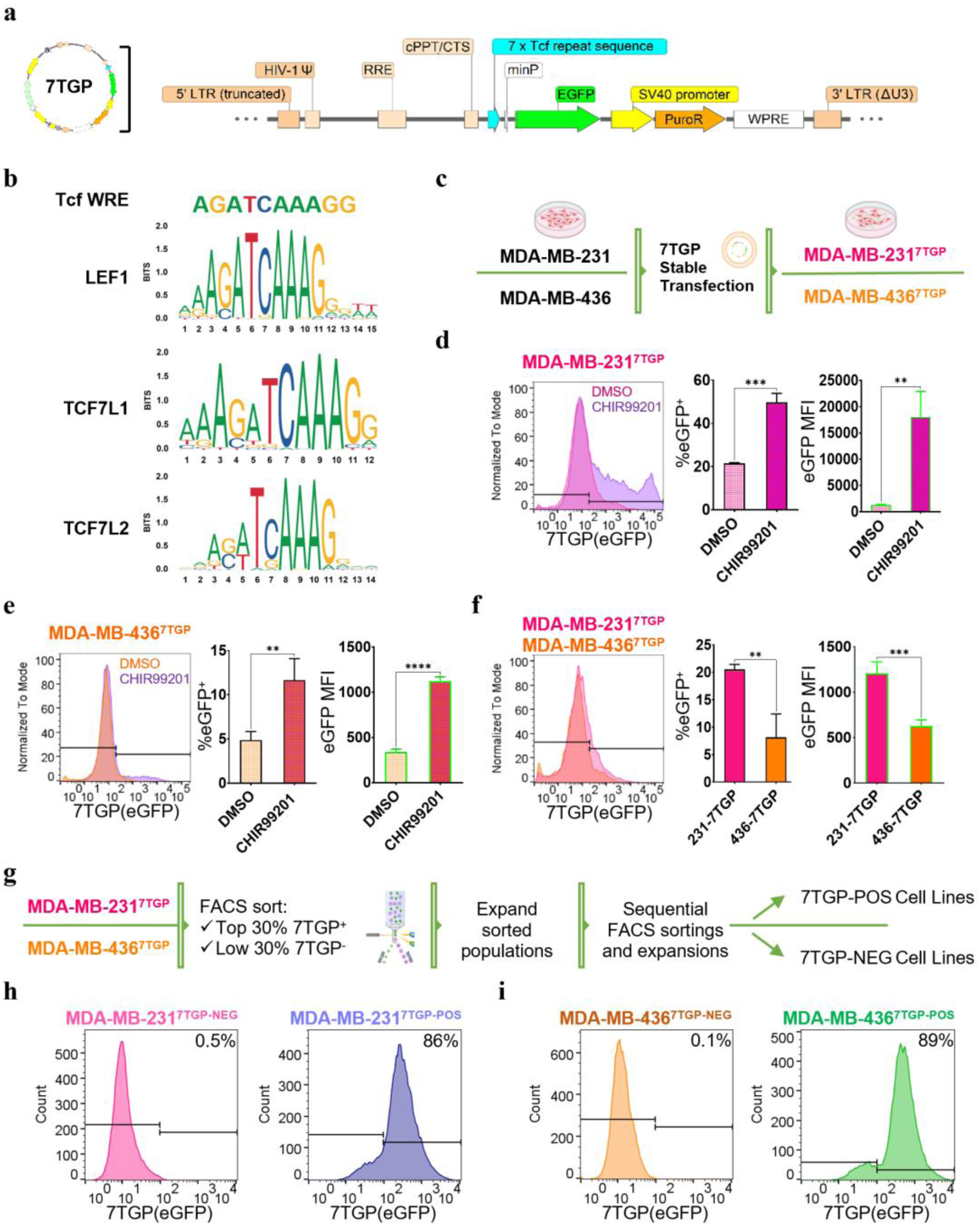
Assessment of Wnt activity and generation of Wnt-positive and Wnt-negative TNBC cell lines using the 7TGP reporter system. **a,** Schematic of the 7TGP Tcf/Lef Wnt activity reporter vector. **b,** DNA sequence of the Wnt Response Element (WRE) within the 7xTcf promoter of the 7TGP vector, with consensus binding sites for LEF1, TCF7L1, and TCF7L2 (logos sourced from JASPAR). **c,** Workflow for generating MDA-MB231^7TGP^ and MDA-MB-436^7TGP^ stable cell lines following puromycin selection. **d, e** Flow cytometry analysis of eGFP fluorescence in MDA-MB-231^7TGP^ (d) and MDA-MB-436^7TGP^ (e) cells treated with DMSO or CHIR99021 (3 µM) for 24 h. Left, eGFP histograms; middle, frequency of eGFP-positive cells; right, mean fluorescence intensity (MFI); n = 2 independent experiments. **f,** Comparative flow cytometry analysis of basal eGFP expression in MDA-MB-231^7TGP^ and MDA-MB-436^7TGP^ cells, shown as histograms (left), frequency (middle), and MFI (right); n = 2. **g,** Schematic of the sorting strategy used to generate Wnt-negative (NEG) and Wnt- positive (POS) sublines based on eGFP expression by FACS. **h, i** Flow cytometry histograms showing eGFP fluorescence intensity in MDA-MB-231^7TGP-NEG^ (pink, h, left), MDA-MB-231^7TGP-POS^ (blue, h, right), MDA-MB-436^7TGP-NEG^ (orange, i, left), and MDA-MB-436^7TGP-POS^ (green, i, right) cell lines. Numbers indicate the percentage of eGFP-positive cells. Error bars represent mean ± s.d. P values were calculated using an unpaired two-tailed t-test (**P < 0.01; ***P < 0.001; **P < 0.0001) in d–f.

To validate this cellular assay, MDA-MB-231^7TGP^ and MDA-MB-436^7TGP^ cells were treated with DMSO or 3 µM CHIR99021, a GSK3β inhibitor known to activate canonical Wnt/β-catenin signaling, for 24 hours. Next, eGFP-positive cell frequency and mean fluorescence intensity (MFI) were assessed by flow cytometry. Treatment with CHIR99021 significantly increased both the frequency of eGFP-positive cells (P = 0.0003) and eGFP MFI (P = 0.004) in MDA-MB-231^7TGP^ cells (Fig. 1d). Similarly, CHIR99021 treatment of MDA-MB-436^7TGP^ cells resulted in a significant increase in the frequency of eGFP-positive cells (P = 0.002) and eGFP MFI (P < 0.0001) (Fig. 1e). These results validate the use of the 7TGP reporter plasmid as a robust tool for evaluating Wnt activity in TNBC cells.

Next, we evaluated the basal Wnt activity in MDA-MB-231^7TGP^ and MDA-MB-436^7TGP^ cells by flow cytometry. Our analysis revealed that the percentage of eGFP-positive cells was 2.5-fold higher in MDA-MB-231^7TGP^ cells compared to MDA-MB-436^7TGP^ cells (P = 0.0013), and the eGFP MFI in MDA-MB-231^7TGP^ cells was 1.9-fold greater than that observed in MDA-MB-436^7TGP^ cells (P = 0.0002) (Fig. 1f).

Together, these results highlight inter- and intra-tumoral heterogeneity in Wnt signaling levels across these mesenchymal TNBC cellular models.

### Establishment of paired Wnt-positive and Wnt-negative TNBC cell lines

Given the observed intra-tumoral heterogeneity in Wnt activity in MDA-MB-231^7TGP^ and MDA-MB-436^7TGP^ cells, we next sought to isolate subpopulations with distinct Tcf/Lef activity statuses. Cells were sorted by FACS based on eGFP fluorescence detected on the FL1 channel, designating the brightest third of eGFP-positive cells as Wnt-positive and the left-most third of eGFP-negative cells as Wnt-negative (Fig. 1g). This sorting procedure was repeated seven times to enrich each population.

From the MDA-MB-231^7TGP^ cell line, we established the MDA-MB-231^7TGP-NEG^ cell line (>99% eGFP-negative) and the MDA-MB-231^7TGP-POS^ cell line (>86% eGFP-positive) (Fig. 1h). Similarly, MDA-MB-436^7TGP-NEG^ and MDA-MB-436^7TGP-POS^ sublines were generated with >99% eGFP-negative cells and >89% eGFP-positive cells, respectively (Fig. 1i). These results demonstrate that subpopulations differing in Wnt activity can be reliably isolated using the 7TGP reporter system, enabling the investigation of Wnt-associated heterogeneity.

### RNA-Seq analysis reveals transcriptomic differences between Wnt-positive and Wnt-negative TNBC cells

Cancer cells adapt to nutrient deprivation by activating transcriptional programs that enhance metabolic plasticity, stress tolerance, and resistance to therapy, all of which contribute to tumor progression. To investigate how Wnt activity intersects with these adaptive responses, we characterized the transcriptomic profiles of Wnt-positive and Wnt-negative TNBC cell lines under both nutrient-limited and nutrient-rich conditions. MDA- MB-231^7TGP-POS^, MDA-MB-231^7TGP-NEG^, MDA-MB-436^7TGP-POS^, and MDA-MB-436^7TGP-NEG^ cells were cultured in low serum (0.5% FBS, 3 days; Supplementary Fig. 1a) or standard serum (10% FBS, 6 days; Supplementary Fig. 1b), followed by total RNA extraction and bulk RNA-Seq for transcriptomic profiling (workflow in Fig. 2a). Examining both low and standard serum conditions enabled us to distinguish Wnt-specific transcriptional programs from nutrient stress responses, assess context-dependent Wnt activity, and identify robust gene expression signatures conserved across metabolic states.

**Figure 2.**
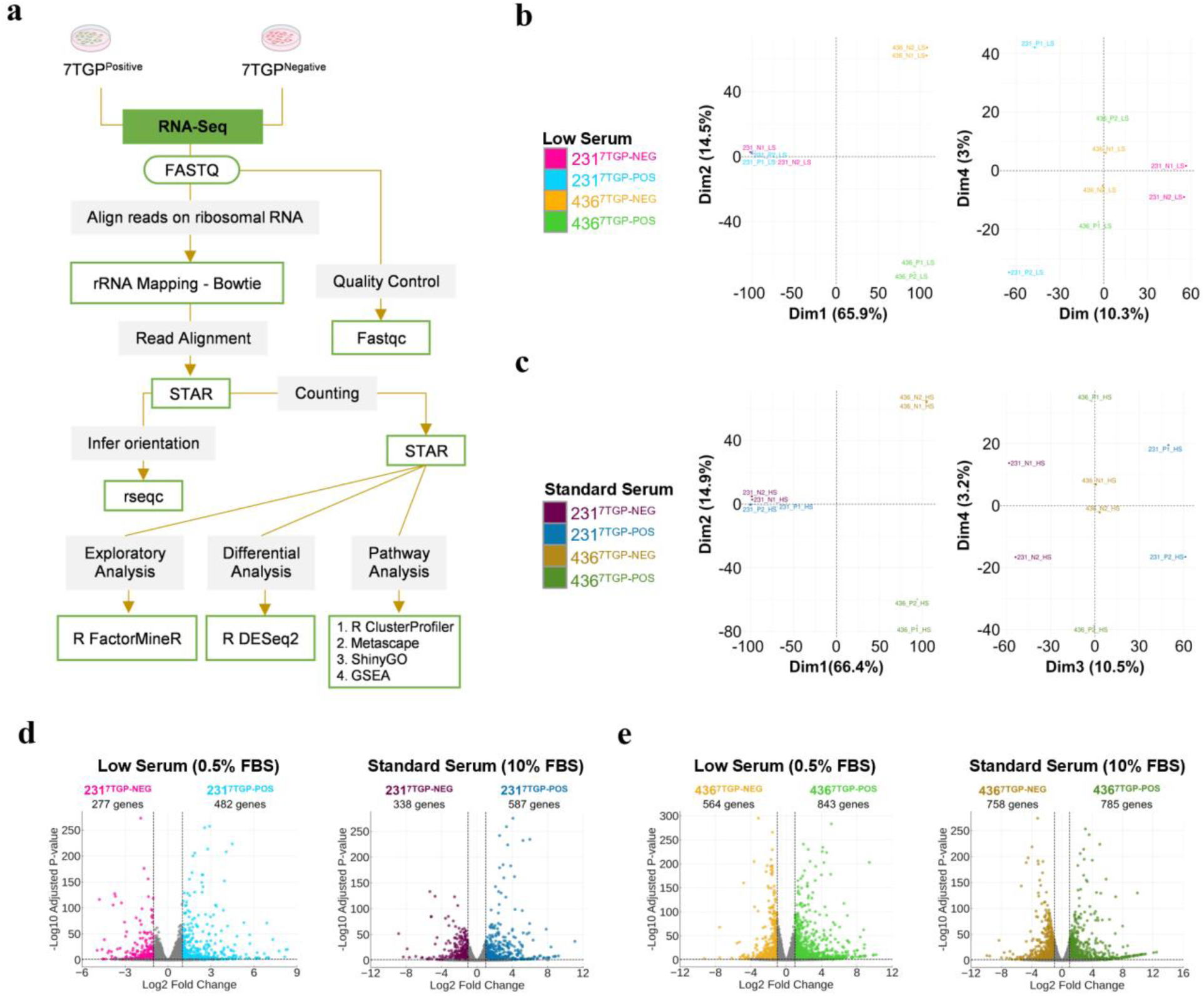
Transcriptomic analysis of Wnt-negative and Wnt-positive TNBC cell lines. a, Schematic representation of the workflow used to profile the transcriptomes of MDA-MB-231^7TGP-NEG^, MDA-MB-231^7TGP-POS^, MDA-MB-436^7TGP-NEG^, and MDA-MB-436^7TGP-POS^ cell lines. **b, c** Principal component analysis (PCA) plots showing sample variance among Wnt-negative and Wnt-positive TNBC cell lines cultured under low serum (b) and standard serum (c) conditions. **d, e** Volcano plots illustrating differential gene expression between MDA-MB-231^7TGP-NEG^ and MDA-MB-231^7TGP-POS^ cells (d) and between MDA-MB- 436^7TGP-NEG^ and MDA-MB-436^7TGP-POS^ cells (e), under low serum (left) and standard serum (right) conditions. Differential expression was defined using an absolute log₂ fold change ≥ 1.

Principal component analysis (PCA) was performed to evaluate variance between samples. Under low serum conditions, 65.9% of the variance was attributed to differences between MDA-MB-231- and MDA-MB-436-derived cell lines, 14.5% to differences between MDA-MB-436^7TGP-POS^ and MDA-MB-436^7TGP-NEG^ cells, 10.3% to differences between MDA-MB-231^7TGP-POS^ and MDA-MB-231^7TGP-NEG^ cells, and less than 3% to variance between biological duplicates (Fig. 2b). Under standard serum conditions, similar patterns were observed: 66.4% variance between MDA-MB-231- and MDA-MB- 436-derived cell lines, 14.9% between MDA-MB-436^7TGP-POS^ and MDA-MB-436^7TGP-NEG^ cells, 10.5% between MDA-MB-231^7TGP-POS^ and MDA-MB-231^7TGP-NEG^ cells, and less than 3.2% between biological duplicates (Fig. 2c). These analyses demonstrate that inter-cell line heterogeneity contributes more significantly to transcriptomic variability than differences in Wnt activity status. Importantly, the low variances (∼3%) observed between biological duplicates under both serum conditions underscore the robustness of our analysis and the capture of biologically meaningful signals.

Differentially expressed genes (DEGs) between Wnt-positive and Wnt-negative cell lines were identified using the DESeq2 R package (v1.26.0). Genes were considered differentially expressed if they exhibited an adjusted P value <0.05 and an absolute log2 fold change (Wnt-positive/Wnt-negative) ≥1, under either low or standard serum conditions.

Between MDA-MB-231^7TGP-POS^ and MDA-MB-231^7TGP-NEG^ cells, 759 DEGs were identified under low serum conditions and 925 DEGs under standard serum conditions (Fig. 2d). Similarly, 1,407 DEGs and 1,543 DEGs were identified between MDA-MB- 436^7TGP-POS^ and MDA-MB-436^7TGP-NEG^ cells under low and standard serum conditions, respectively (Fig. 2e). In all comparisons, DEGs included both genes upregulated and downregulated when comparing each 7TGP-positive cell line to its corresponding 7TGP- negative counterpart. The complete lists of DEGs for each condition are provided in Supplementary File 1. The top 100 DEGs between MDA-MB-231^7TGP-POS^ and MDA-MB- 231^7TGP-NEG^ cells under low and standard serum conditions are depicted as heatmaps in Supplementary Fig. 2 and 3 (left panels), respectively. Similarly, the top 100 DEGs between MDA-MB-436^7TGP-POS^ and MDA-MB-436^7TGP-NEG^ cells are depicted as heatmaps under low and standard serum conditions in Supplementary Fig. 2 and 3 (right panels), respectively. Genes previously annotated as Wnt-related genes (Supplementary File 2) are marked in black along the y-axis of each heatmap.

Collectively, these findings demonstrate that Wnt-positive and Wnt-negative TNBC cell lines exhibit distinct transcriptomic profiles, validating the impact of Wnt activity on gene expression and enabling downstream pathway analyses.

### GO Biological Process analysis: Wnt activity enriches immune, developmental, and ECM terms under low and standard serum in TNBC cells

To explore the biological significance of DEGs between Wnt-positive and Wnt-negative TNBC cells, Gene Ontology Biological Process (GO BP) enrichment analyses were conducted using clusterProfiler v3.14.0. DEGs (upregulated and downregulated in Wnt-positive versus Wnt-negative) with an absolute log₂ fold change ≥ 1 were included, and GO terms with an adjusted P value < 0.05 were considered significant.

For GO BP analysis, DEGs between MDA-MB-231^7TGP-POS^ and MDA-MB-231^7TGP- NEG^ cells revealed significant enrichment of 473 and 367 terms under low and standard serum conditions, respectively, with 236 shared between conditions (Fig. 3a, left panels). Among the top shared terms were processes related to immune and inflammatory signaling (e.g., response to lipopolysaccharide, myeloid leukocyte migration), as well as developmental regulation (e.g., nervous system development, epithelial cell proliferation), suggesting consistent activation of immune and developmental transcriptional programs. Unique enrichments under low serum included mononuclear cell migration and second-messenger-mediated signaling, while standard serum-specific terms featured processes such as homophilic cell adhesion and SMAD protein phosphorylation (Fig. 3a, right panel). A full list of enriched terms is available in Supplementary File 3.

**Figure 3.**
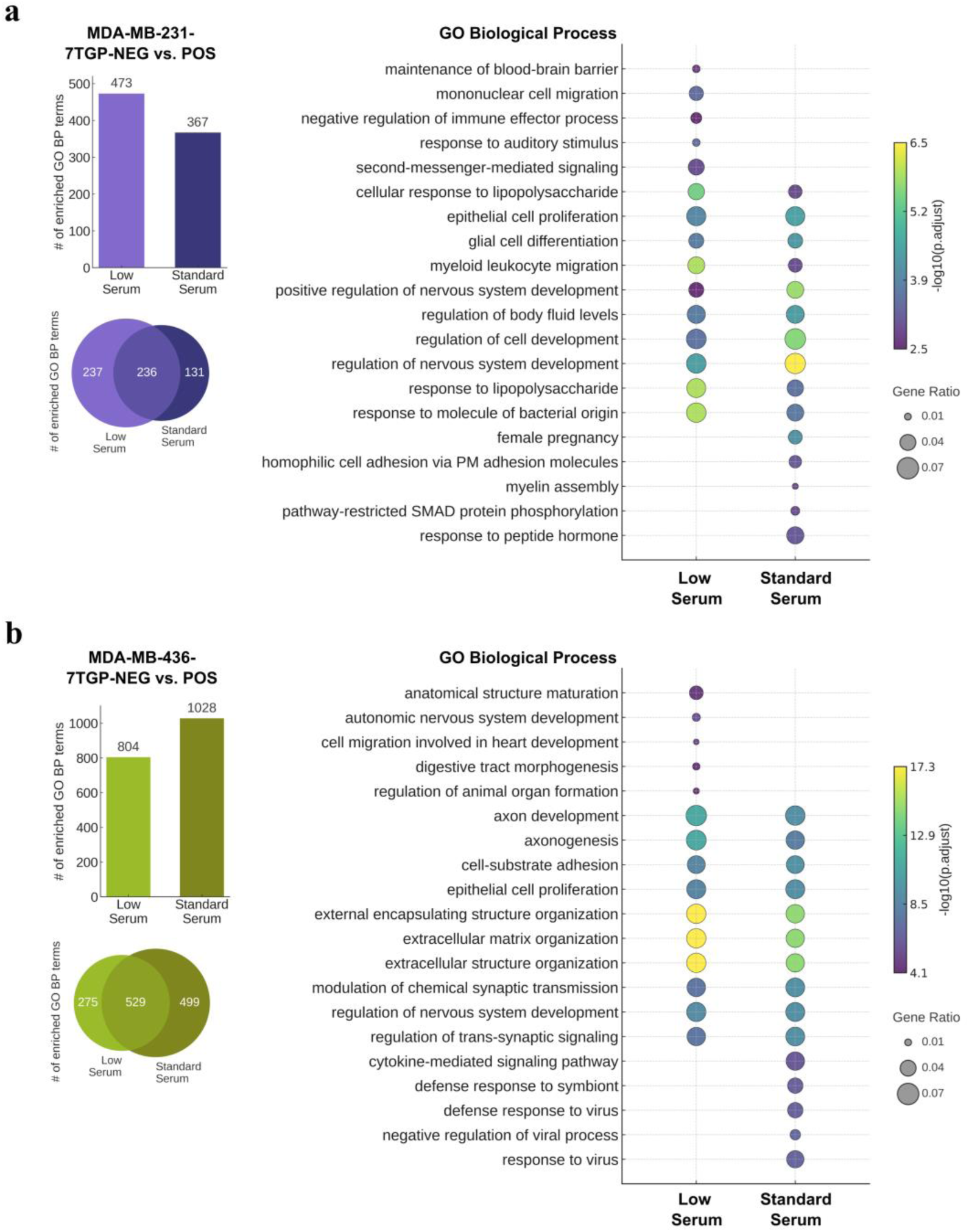
Gene Ontology Biological Process analyses of differentially expressed genes between Wnt-negative and Wnt-positive TNBC cell lines. **a**, GO Biological Process (GO BP) terms significantly enriched (adjusted P < 0.05) among DEGs (log₂ fold change ≥ 1) between MDA-MB-231^7TGP-NEG^ and MDA-MB-231^7TGP-POS^ cultured in low or standard serum. Bar graph (top left) shows the number of enriched terms per condition; Venn diagram (bottom left) illustrates the overlap. Bubble plot (right) presents the top five condition-specific GO BP terms ranked by adjusted P value, and the top ten shared terms selected using Stouffer’s method for combined adjusted P values. **b**, GO BP terms significantly enriched (adjusted P < 0.05) among DEGs (log₂ fold change ≥ 1) between MDA-MB-436^7TGP-NEG^ and MDA-MB-436^7TGP-POS^ under low or standard serum. Bar graph (top left) and Venn diagram (bottom left) show the number and overlap of enriched terms. Bubble plot (right) presents the top five unique terms ranked by adjusted P value and the top ten shared terms selected using Stouffer’s method.

For DEGs between MDA-MB-436^7TGP-POS^ and MDA-MB-436^7TGP-NEG^ cells, 804 and 1,028 GO BP terms were enriched under low and standard serum conditions, respectively, with 529 shared between conditions (Fig. 3b, left panels).

Shared enrichment patterns included processes related to structural organization and neurodevelopment, such as extracellular matrix (ECM) organization, axonogenesis, and regulation of nervous system development, highlighting biological programs associated with differential Wnt activity. Under low serum, uniquely enriched terms emphasized developmental and morphogenic processes (e.g., digestive tract morphogenesis, autonomic nervous system development), while standard serum-specific enrichments were characterized by antiviral and cytokine signaling pathways (e.g., defense response to virus, cytokine-mediated signaling pathway) (Fig. 3b, right panel). The full list of enriched terms is provided in Supplementary File 3.

Although both comparisons between Wnt-negative and Wnt-positive cells showed shared enrichment in developmental and immune-related GO BP terms, differentially expressed genes between MDA-MB-231^7TGP-NEG^ and MDA-MB-231^7TGP-POS^ cells under low serum were more strongly associated with immune signaling pathways. In contrast, differentially expressed genes between MDA-MB-436^7TGP-NEG^ and MDA-MB-436^7TGP-POS^ cells were enriched for morphogenesis and ECM organization. Under standard serum, immune-related pathways emerged more prominently in the MDA-MB-436-7TGP comparison.

Together, these findings offer mechanistic insight into the transcriptional programs modulated by Wnt signaling in TNBC cells under varying nutrient conditions. The combined GO BP enrichment results provide a functional framework for understanding how Wnt activity shapes immune signaling, developmental programming, matrix remodeling, and receptor-mediated communication in a cellular context-dependent manner.

### GO Molecular Function analysis: serum-specific enrichment of ECM, receptor, and ion transport functions linked to Wnt activity in TNBC cells

To investigate how Wnt activity shapes molecular functions under distinct nutrient conditions, we performed GO Molecular Function (GO MF) enrichment analysis on DEGs from each Wnt-positive TNBC cell line, relative to its Wnt-negative counterpart, under low and standard serum conditions. We then examined the functional overlap between MDA- MB-231^7TGP-POS^ and MDA-MB-436^7TGP-POS^ cells within each condition to identify molecular functions consistently associated with Wnt activity.

MDA-MB-231^7TGP-POS^ and MDA-MB-436^7TGP-POS^ cells were cultured in low or standard serum conditions, and differential expression analysis was performed to compare their transcriptomic profiles with those of their respective 7TGP-negative counterparts (1 log₂ fold-change cutoff). Under low serum conditions, 759 DEGs were identified in the MDA-MB-231^7TGP-POS^ cell line, and 1407 DEGs were identified in the MDA-MB-436^7TGP-POS^ cell line (Circos Plot, Fig. 4a). Across the two 7TGP-positive cell lines cultured in low serum, 216 genes overlapped (Circos plot, Fig. 4a), with 138 consistently regulated (105 up, 33 down) and 78 showing opposite trends (Upset Plot, Fig. 4a).

**Fig. 4.**
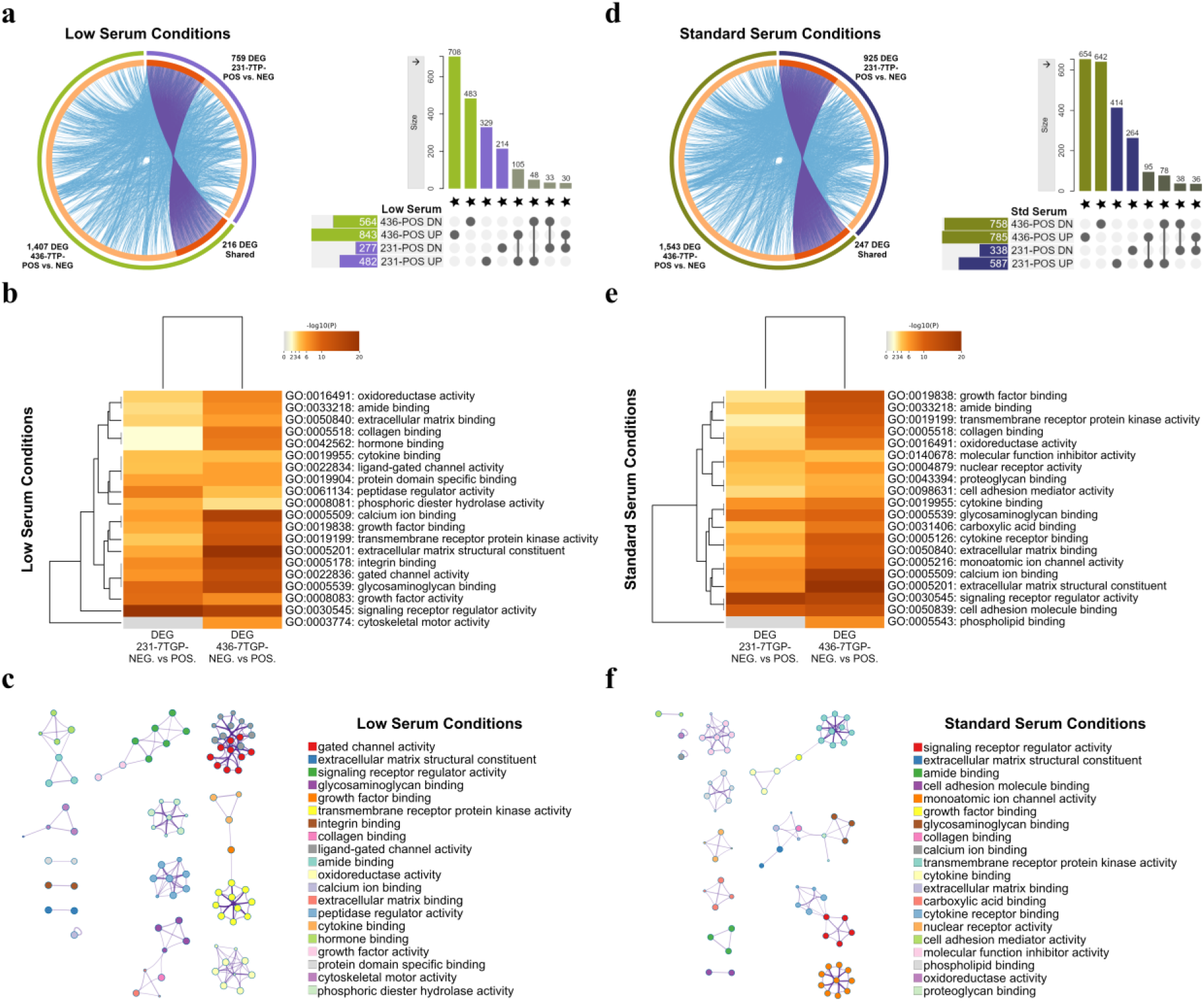
Overlap and functional enrichment analysis of Wnt-regulated gene signatures in TNBC cells under low and standard serum conditions. a–c. DEGs identified between Wnt-positive and Wnt-negative TNBC cell lines cultured under low serum (0.5% FBS) conditions. **a, left** Circos plot generated by Metascape showing overlap between DEGs from MDA-MB-231^7TGP-POS^ versus MDA-MB-231^7TGP-NEG^ (759 DEGs) and MDA-MB-436^7TGP-POS^ vs MDA-MB-436^7TGP-NEG^ (1,407 DEGs). Purple links indicate shared 216 DEGs; blue connections represent known interactions. **a, right** UpSet plot illustrating intersections among upregulated and downregulated DEGs from MDA-MB-231^7TGP-POS^ (482 up, 277 down) and MDA-MB-436^7TGP-POS^ (843 up, 564 down) comparisons. **b,** Heat map of the top 20 enriched GO MF categories identified by Metascape analysis, clustered by Kappa- statistical similarity and colored by P value. **c,** Network visualization of enriched GO MF categories from Metascape analysis, with node size proportional to gene number and node color reflecting cluster identity. **d–f** DEGs identified between Wnt-positive and Wnt-negative TNBC cell lines cultured under standard serum (10% FBS) conditions. **d, left** Circos plot generated by Metascape showing overlap between DEGs from MDA-MB-231^7TGP-POS^ versus MDA-MB-231^7TGP-NEG^ (925 DEGs) and MDA-MB-436^7TGP-POS^ versus MDA-MB-436^7TGP-NEG^ (1,543 DEGs). Purple links indicate shared 247 DEGs; blue connections represent known interactions. **d, right** UpSet plot illustrating intersections among upregulated and downregulated DEGs from MDA-MB-231^7TGP-POS^ (587 up, 388 down) and MDA-MB-436^7TGP-POS^ (785 up, 758 down) comparisons. **e,** Heat map of the top 20 enriched GO MF categories identified by Metascape analysis under standard serum conditions. **f,** Network visualization of enriched GO MF categories from Metascape analysis, with node size indicating gene number and node color reflecting either cluster identity.

To characterize the functional landscape of these DEGs, GO MF analysis was performed using Metascape. This approach selects the most significant terms within each cluster of related annotations, allowing for broader interpretation of functional patterns. The top 20 GO MF clusters derived from DEGs in MDA-MB-231^7TGP-POS^ and MDA-MB-436^7TGP-POS^ cells under low serum conditions are displayed in a P value–ranked heatmap (Fig. 4b), and the complete list of 138 terms encompassing the top 20 cluster groups is provided in Supplementary File 4. A corresponding network diagram visualizes the cluster relationships and associated significance levels (Fig. 4c). Among these, the 10 most recurrent genes in the enriched GO MF categories were: *FGFR2*, *EPHA7*, *ALK*, *KIT*, *NTRK1*, *PDGFRB*, *FGFR3*, *FLT1*, *ERBB4*, and *NTRK2* all receptor tyrosine kinases (Supplementary File 4).

Under low serum conditions, DEGs from MDA-MB-231^7TGP-POS^ and MDA-MB-436^7TGP-POS^ cells were enriched for GO MF terms related to ECM organization, adhesion, and ligand– receptor signaling (Fig. 4b). Shared terms included collagen binding, glycosaminoglycan binding, and signaling receptor regulator activity, alongside additional enrichments in enzymatic functions, ion transport, and protein binding. Cytoskeletal motor activity was unique to MDA-MB-436^7TGP-POS^ cells.

Among the top-ranked terms, MDA-MB-231^7TGP-POS^ DEGs were most associated with signaling receptor regulator activity, glycosaminoglycan binding, and growth factor activity, while MDA-MB-436^7TGP-POS^ DEGs were enriched for ECM structural constituent, calcium ion binding, and signaling receptor regulator activity.

To assess transcriptional responses under nutrient-rich conditions, we also examined DEGs in both cell lines cultured in standard serum. In the MDA-MB-231^7TGP- POS^ cell line, 925 DEGs were differentially expressed, while the MDA-MB-436^7TGP-POS^ cell line showed 1,543 DEGs expressed, compared to their 7TGP-negative counterparts (Circos Plot, Fig. 4d). Across the two lines, 247 DEGs overlapped (Circos plot, Fig. 4d), with 133 consistently regulated (95 up, 38 down) and 114 showing opposite trends (Upset Plot, Fig. 4d).

The top 20 GO MF clusters enriched by DEGs of the MDA-MB-231^7TGP-POS^ and MDA-MB-231^7TGP-POS^ cell lines cultured in standard serum are shown in a P-value–ranked heatmap (Fig. 4e), with the full list of 105 terms encompassing the top 20 cluster groups in Supplementary File 4. The top 20 enriched GO MF terms for the standard serum conditions are also displayed in a network diagram to visualize the cluster relationships and associated significance levels (Fig. 4f). The 10 most frequent genes in the enriched GO MF terms for the differentially expressed genes under standard serum conditions were: *EPHA7*, *ALK*, *KDR*, *KIT*, *FLT4*, *NTRK2*, *PDGFRB*, *RET*, *AXL*, and *FLT1*, which are also receptor tyrosine kinases involved in growth factor, angiogenic, and neurotrophic signaling (Supplementary File 4).

Under standard serum conditions, GO MF enrichment analysis of DEGs from MDA-MB-231^7TGP-POS^ and MDA-MB-436^7TGP-POS^ cells highlighted functions related to ECM organization, ligand–receptor signaling, and molecular binding (Fig. 4f). Shared enrichments included ECM structural constituent, collagen and glycosaminoglycan binding, growth factor binding, and signaling receptor regulator activity. Additional terms included phospholipid binding (specific to MDA-MB-436^7TGP-POS^), oxidoreductase activity, and calcium ion binding. Among the top-ranked terms, DEGs from MDA-MB-231^7TGP-POS^ cells were most enriched for signaling receptor regulator activity, cell adhesion molecule binding, and glycosaminoglycan binding, whereas MDA-MB-436^7TGP-POS^ cells showed strongest enrichment for ECM structural constituent, calcium ion binding, and signaling receptor regulator activity.

Comparison of GO MF profiles across serum conditions revealed both shared and distinct features. Signaling receptor regulator activity ranked highly in both cell lines and conditions, indicating a common reliance on receptor-mediated signaling. Low serum favored matrix remodeling and enzymatic functions, while standard serum emphasized ligand binding, ion transport, and adhesion, reflecting nutrient-dependent transcriptional adaptations.

### KEGG analysis: nutrient-dependent immune and oncogenic pathways associated with Wnt activity in TNBC cells

To investigate which signaling pathways may be modulated by Wnt activation in TNBC cells, KEGG pathway enrichment analyses were performed using the ShinyGO v0.82 bioinformatics tool. Genes differentially expressed in each Wnt-positive cell line relative to its Wnt-negative counterpart, with an absolute log₂ fold change ≥ 1 and a false discovery rate (FDR) < 0.05, were included.

Under low serum conditions, 21 and 41 KEGG pathways were enriched in MDA- MB-231^7TGP-POS^ and MDA-MB-436^7TGP-POS^ cells, respectively, with 8 shared pathways, including cytokine–cytokine receptor interaction, PI3K-Akt signaling, and Wnt signaling (Supplementary File 5). These pointed to common Wnt-associated activation of immune, ligand–receptor, and oncogenic pathways under nutrient-limited conditions. Under standard serum, 25 pathways were enriched in MDA-MB-231^7TGP-POS^ cells and 52 in MDA-MB-436^7TGP-POS^ cells, with 19 shared, including pro-inflammatory signaling (e.g., IL- 17, TNF, JAK-STAT, NF-κB) and cancer-related pathways. Additional overlaps in axon guidance, ECM-receptor interaction, and focal adhesion suggest coordinated regulation of migration, matrix remodeling, and cytoskeletal dynamics in nutrient-rich environments. Notably, the Wnt signaling pathway (KEGG) was enriched under low serum conditions by DEGs from MDA-MB-231^7TGP-POS^ versus MDA-MB-231^7TGP-NEG^, and MDA- MB-436^7TGP-POS^ versus MDA-MB-436^7TGP-NEG^ (pathway enrichment shown in Supplementary Fig. 4 and 5, respectively), but not under standard serum conditions, suggesting the emergence of a distinct subset of Wnt-related genes in serum-rich environments.

Together, these findings support the role of Wnt activation in regulating diverse signaling pathways relevant to tumor progression, immune modulation, and extracellular communication in TNBC cells.

### Gene Set Enrichment Analysis: contrasting enrichment of EMT, inflammatory, and metabolic gene programs defined by Wnt activity in TNBC cells

To explore how Wnt signaling aligns with established transcriptomic programs, we performed Gene Set Enrichment Analysis (GSEA) using Hallmark gene sets from the Molecular Signatures Database (MSigDB). The analysis was conducted on all detected genes ranked by log₂ fold change (Supplementary File 6), and gene sets were considered significantly enriched at P < 0.05 and FDR < 0.25 (Supplementary File 7).

In the comparison between MDA-MB-231^7TGP-POS^ and MDA-MB-231^7TGP-NEG^ cells, GSEA identified gene sets differentially enriched based on Wnt activity under both serum conditions. Gene sets enriched at the top of the ranked list correspond to higher expression in MDA-MB-231^7TGP-POS^ cells, while those enriched at the bottom indicate higher expression in MDA-MB-231^7TGP-NEG^ cells. Under low serum, the expression profile of MDA-MB-231^7TGP-NEG^ cells was enriched for *cholesterol homeostasis* (Fig. 5a), while under standard serum, these cells showed enrichment of *MYC targets V2* and *estrogen response late* gene sets (Fig. 5b). In contrast, the expression profile of MDA-MB-231^7TGP- POS^ cells exhibited enrichment of *Epithelial-to-Mesenchymal Transition (EMT), KRAS signaling up*, *inflammatory response*, *TNFα signaling via NF-κB*, and *IL6/JAK/STAT3 signaling* across both serum conditions. Additional enrichment under standard serum in MDA-MB-231^7TGP-POS^ cells included *interferon-α response*, *interferon-γ response*, *complement*, and *allograft rejection* gene sets (Fig. 5b), indicating context-specific activation of immune and inflammatory pathways in the presence of Wnt activity.

**Fig. 5.**
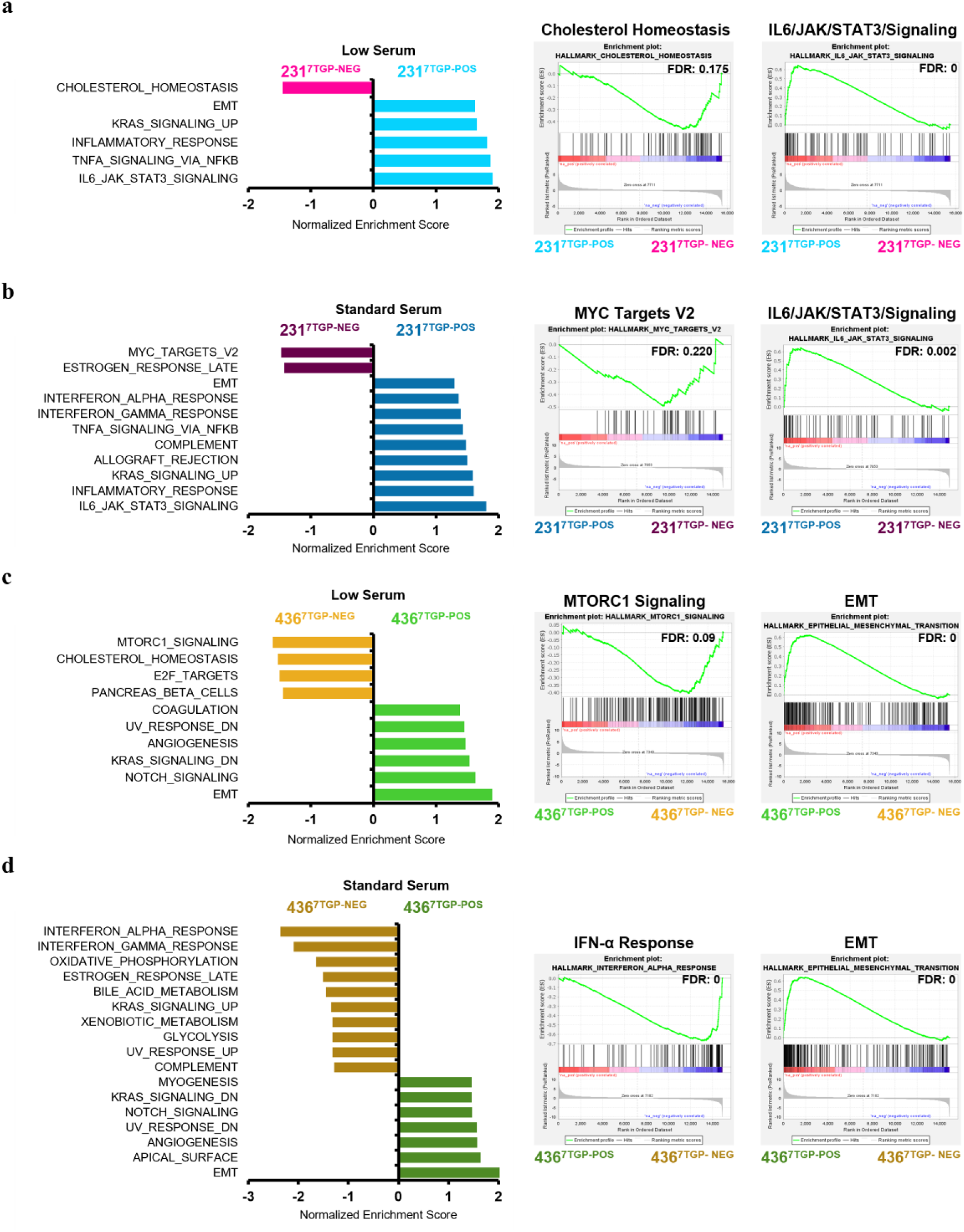
Gene Set Enrichment Analysis (GSEA) of transcriptional programs associated with Wnt activity in TNBC cell lines. GSEA was performed using all detected genes ranked by log₂ fold change between Wnt-negative and Wnt-positive TNBC cells, using MSigDB Hallmark gene sets (P value < 0.05 and FDR < 0.25). **a, b** GSEA results for MDA-MB-231^7TGP-NEG^ and MDA-MB-231^7TGP-POS^ cells cultured under low serum (a) or standard serum (b) conditions. **c, d** GSEA results for MDA-MB-436^7TGP-NEG^ and MDA-MB- 436^7TGP-POS^ cells cultured under low serum (c) or standard serum (d) conditions. Panels display the enriched Hallmark gene sets (left), the top enriched gene set in Wnt-negative cells (middle), and the top enriched gene set in Wnt-positive cells (right).

GSEA was also performed on all detected genes ranked by log₂ fold change from the comparison of MDA-MB-436^7TGP-POS^ and MDA-MB-436^7TGP-NEG^ cells. Similarly, gene sets enriched at the top of the ranked list indicate higher expression in MDA-MB-436^7TGP- POS^ cells, whereas gene sets enriched at the bottom reflect higher expression in MDA- MB-436^7TGP-NEG^ cells. Under low serum conditions, the expression profile of MDA-MB- 436^7TGP-NEG^ cells was enriched for *mTORC1 signaling*, *cholesterol homeostasis*, *E2F targets*, and *pancreas beta cells* (Fig. 5c). Under standard serum, additional gene sets enriched in MDA-MB-436^7TGP-NEG^ cells included *interferon-α response*, *interferon-γ response*, *oxidative phosphorylation*, *estrogen response (late)*, *bile acid metabolism*, *KRAS signaling up*, *xenobiotic metabolism*, *glycolysis*, *UV response up*, and *complement* (Fig. 5d). In contrast, the expression profile of MDA-MB-436^7TGP-POS^ cells showed consistent enrichment of *UV response down*, *angiogenesis*, *KRAS signaling down*, *Notch signaling*, and *EMT* under both serum conditions. Additional enrichment in MDA-MB- 436^7TGP-POS^ cells included *coagulation* under low serum and *myogenesis* and *apical surface* under standard serum (Fig. 5c–d).

Together, these results demonstrate that 7TGP-positive TNBC cell lines consistently activate transcriptional programs associated with EMT, inflammatory signaling, and KRAS-related pathways. In contrast, their 7TGP-negative counterparts tend to enrich pathways related to metabolic regulation, cholesterol homeostasis, cell proliferation, and immune responses, with broader pathway diversity observed in MDA- MB-436^7TGP-NEG^ cells. These findings underscore that Wnt activity delineates distinct transcriptional profiles in TNBC cell lines.

### A Wnt-driven 55-gene expression signature highlights EMT, immune signaling, and ECM functions in TNBC cells

To define a shared transcriptional signature associated with Wnt activity in TNBC models, we identified 42 genes consistently upregulated and 13 genes consistently downregulated in both MDA-MB-231^7TGP-POS^ and MDA-MB-436^7TGP-POS^ cell lines relative to their Wnt-negative counterparts under low and standard serum conditions (Fig. 6a).

**Fig. 6.**
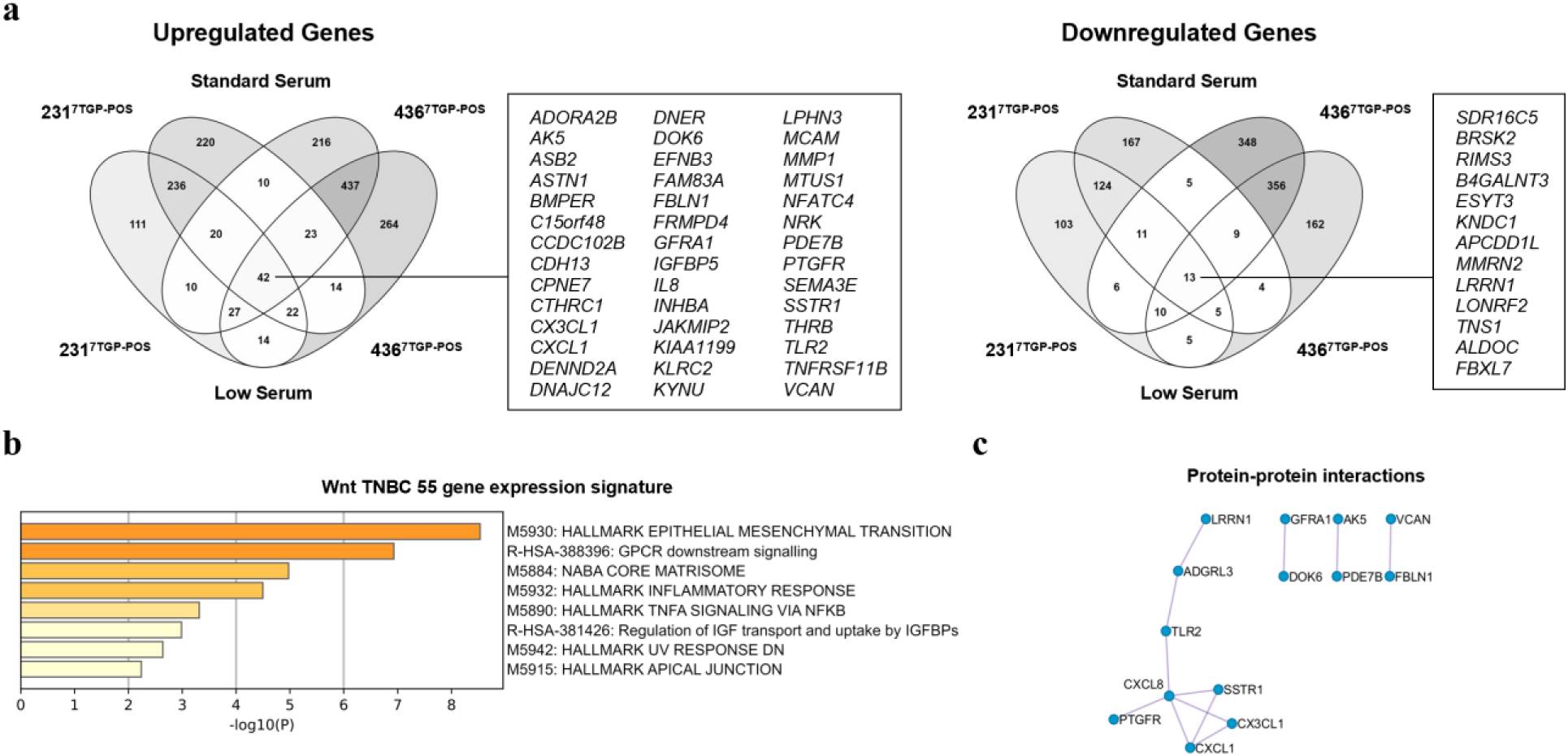
Shared Wnt-regulated transcriptional programs across TNBC cell lines and serum conditions. a, Venn diagrams showing the overlap of DEGs in MDA-MB-231^7TGP-POS^ and MDA-MB-436^7TGP- POS^ cells cultured under low or standard serum conditions. Left: upregulated DEGs; right: downregulated DEGs. A total of 55 DEGs (42 upregulated, 13 downregulated) were shared across both cell lines and serum conditions. **b,** Metascape enrichment analysis of Hallmark and Reactome pathway terms derived from the 55 shared DEGs (–log₁₀(*P*) values). **c,** Protein–protein interaction (PPI) network of the 55 shared DEGs constructed using integrated PPI databases in Metascape.

To assess whether this 55-gene expression signature recapitulated the pathway-level trends identified in the broader DEG datasets, we performed Metascape enrichment analysis using the MSigDB Hallmark and Reactome databases (Fig. 6b). This analysis identified five significantly enriched Hallmark pathways (*EMT*, *NABA core matrisome, inflammatory response, TNF*α *signaling via NF-κB*, *UV response down*, and *apical junction*) and two significantly enriched Reactome pathways (*GPCR downstream signaling* and *regulation of IGF transport and uptake by IGFBPs*). Among these, EMT showed the strongest enrichment, highlighting its association with Wnt-driven mesenchymal programs in TNBC. In addition, several enriched pathways, including inflammatory response, TNFα signaling via NF-κB, and GPCR downstream signaling, reflected the involvement of inflammatory and immune-related processes.

Next, we performed protein–protein interaction (PPI) analysis of the 55-gene expression signature using Metascape (Fig. 6c). MCODE clustering identified four enriched modules, with the largest cluster comprising chemokines, immune-related receptors, and GPCR- and receptor-associated proteins, including LRRN, ADGRL3, TLR2, CXCL8, SSTR1, PTGFR, CXCL1, and CX3CL1 (Fig. 6c).

Together, these findings demonstrate that the 55-gene expression signature captures Wnt-driven transcriptional programs linked to mesenchymal transition, immune modulation, ECM remodeling, growth factor signaling, cellular stress responses, and epithelial organization, key processes implicated in TNBC progression.

### Wnt-related gene expression signatures from TNBC cell lines are preferentially enriched in MLIA TNBC tumors

To evaluate the clinical relevance of the Wnt-related transcriptional programs identified in TNBC cell lines, we next investigated their expression patterns in primary tumor samples from TNBC patients. We assessed the expression of four Wnt-related gene sets derived from differential expression analysis between Wnt-positive and Wnt-negative TNBC cell lines. GES1 consists of DEGs between MDA-MB-231^7TGP-POS^ and MDA-MB-231^7TGP-NEG^ cells cultured in low serum; GES2 consists of DEGs between MDA- MB-436^7TGP-POS^ and MDA-MB-436^7TGP-NEG^ cells cultured in low serum; GES3 consists of DEGs between MDA-MB-231^7TGP-POS^ and MDA-MB-231^7TGP-NEG^ cells cultured in standard serum; and GES4 consists of DEGs between MDA-MB-436^7TGP-POS^ and MDA-MB- 436^7TGP-NEG^ cells cultured in standard serum. Expression of each GES was assayed in a TNBC RNA-Seq cohort (Supplementary File 8).

Among the different TNBC subtypes (LAR [n = 137], MLIA [n = 121], BLIA [n = 199], and BLIS [n = 242]), MLIA tumors consistently exhibited significantly higher expression scores for GES1, GES2, and GES3 compared to LAR, BLIA, and BLIS (Fig. 7a–c). For GES4, MLIA tumors showed significantly elevated scores relative to LAR and BLIA, but not BLIS (Fig. 7d). Correlation analysis between the four gene sets and the Wnt/β-catenin signature (WBCS) defined by Lehmann *et al*.^6^ revealed the strongest correlation for GES2 (r = 0.79), followed by GES3 (r = 0.57), GES1 (r = 0.56), and GES4 (r = 0.45) (Supplementary Fig. 8). Together, these results demonstrate that human MLIA tumors most strongly reflect the Wnt-related transcriptional programs identified in TNBC cell lines. Moreover, the moderate correlations between GES1–GES4 and the canonical Wnt/β-catenin signature suggest that these gene sets may capture Wnt-linked but distinct transcriptional programs, potentially reflecting novel aspects of Wnt signaling in TNBC.

**Fig. 7.**
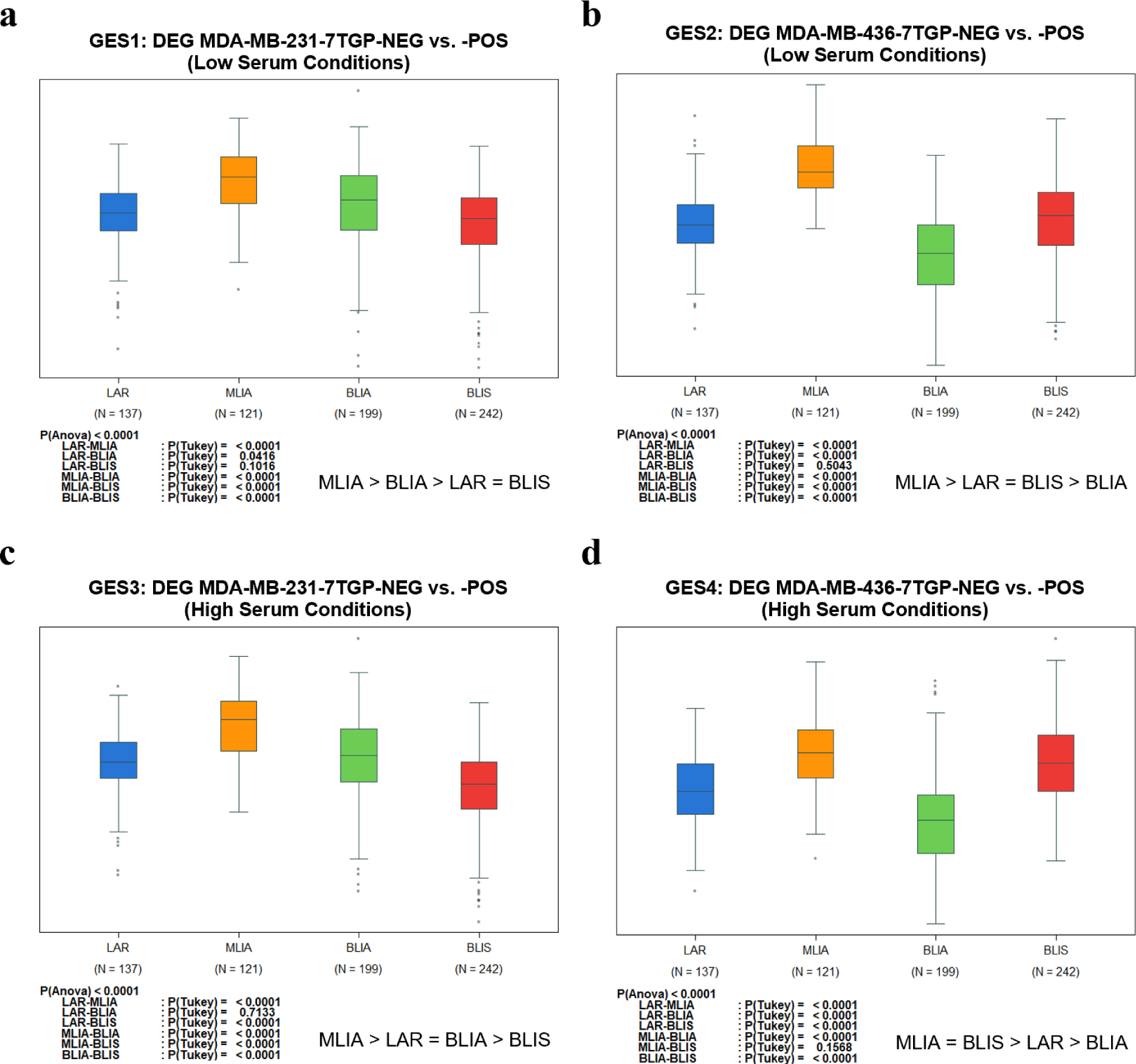
Expression of Wnt-related gene signatures across human TNBC subtypes. Four Wnt-related gene expression signatures (GES1–GES4), derived from differential expression analysis of Wnt-positive and Wnt-negative TNBC cell lines, were evaluated in a cohort of TNBC patient tumors (n = 699). The cohort includes four molecular TNBC subtypes: LAR, MLIA, BLIA, and BLIS. Bar plots show signature scores across subtypes for each gene expression signature: **a,** GES1: 687 DEGs between MDA-MB-231^7TGP-POS^ and MDA-MB-231^7TGP-NEG^ cells cultured in low serum. **b,** GES2: 1,278 DEGs between MDA-MB-436^7TGP- POS^ and MDA-MB-436^7TGP-NEG^ cells cultured in low serum. **c,** GES3: 848 DEGs between MDA-MB-231^7TGP- POS^ and MDA-MB-231^7TGP-NEG^ cells cultured in standard serum. **d,** GES4: 1,418 DEGs between MDA-MB- 436^7TGP-POS^ and MDA-MB-436^7TGP-NEG^ cells cultured in standard serum. Statistical significance was assessed using one-way ANOVA with Tukey’s post hoc test. P values ≤ 0.01 were considered significant and are shown below each panel.

Heatmap visualization and hierarchical clustering of each gene set (GES1–GES4) across the TNBC RNA-Seq cohort identified subsets of genes with distinct expression patterns across TNBC subtypes (Supplementary Fig. 9). These primary clusters, referred to as H1 clusters, contained 87 genes for GES1, 56 for GES2, 72 for GES3, and 119 for GES4. An additional cluster was identified for GES3, containing 204 genes and designated H2 (Supplementary File 9). When comparing the expression of the H1 clusters across the TNBC subtypes, MLIA consistently exhibited the highest expression (Supplementary Fig. 10). Correlation analysis with the WBCS revealed strong correlations for H1 GES1 (r = 0.86), H1 GES4 (r = 0.85), H2 GES3 (r = 0.81), and H1 GES3 (r = 0.81), whereas H1 GES2 showed a weaker correlation (r = 0.57) (Supplementary Fig. 11). Inter-cluster comparisons showed that H2 GES3 was highly correlated with H1 GES1 (r = 0.94), H1 GES3 (r = 0.95), and H1 GES4 (r = 0.93) (Supplementary Fig. 11).

GO BP enrichment analysis of the H1 clusters (and H2 for GES3) revealed six core biological processes consistently enriched across all four gene sets: *vasculature development*, *circulatory system development*, *locomotion*, *cell migration*, *regulation of locomotion*, and *regulation of cell migration* (Supplementary File 9). These clusters represent transcriptionally active subsets within each Wnt-related GES, originally derived from TNBC cell lines, that most strongly capture subtype-associated variation across the tumor cohort. Their consistent enrichment in MLIA tumors underscores their potential as core Wnt-responsive modules capable of distinguishing MLIA from other TNBC subtypes. Wnt-associated GES from TNBC cell lines were preferentially expressed in MLIA tumors, indicating subtype-specific activation of Wnt-responsive programs. Clustering identified core subsets (H1/H2) most strongly aligned with MLIA and the canonical Wnt/β- catenin signature. GO BP analysis of these clusters revealed consistent enrichment for migratory and vascular processes, supporting their relevance as functional markers of Wnt activity in MLIA.

### Tukey’s honestly significant difference test analysis: Wnt-related gene sets distinguish MLIA from other TNBC subtypes

Given the clinical and molecular heterogeneity of TNBC, we assessed whether Wnt-related gene sets could stratify tumors by subtype using a quantitative variance- based approach. We applied Tukey’s Honestly Significant Difference (HSD) analysis to eight gene sets (Table 1), derived from GES1–GES4 and subdivided based on upregulation or downregulation in 7TGP-positive versus 7TGP-negative cells.

**Table 1.** Discriminatory power of Wnt-related gene signatures across TNBC subtypes. Tukey’s Honestly Significant Difference (HSD) statistic was used to evaluate the ability of Wnt-related gene sets to distinguish the MLIA subtype (n = 121) from other TNBC subtypes (LAR [n = 137], BLIA [n = 199], and BLIS [n = 242]). For each gene set, the minimum HSD value across the three pairwise comparisons is reported. Gene sets with higher minimum HSD values were considered more discriminatory, and the top-performing gene set was selected as the optimal signature.

| **Gene Expression Signature** | **Serum Condition** | **HSD**<br>**MLIA-LAR** | **HSD**<br>**MLIA-BLIA** | **HSD**<br>**MLIA-BLIS** | **HSD**<br>**Minimum** |
| --- | --- | --- | --- | --- | --- |
| 1_MDA-MB-231<sup>7TGP-POS</sup> Upregulated | Low | 9.296 | 12.0883 | 12.4921 | 9.296 |
| 2_MDA-MB-231<sup>7TGP-POS</sup> Downregulated | Low | 7.4858 | 15.8397 | 9.2341 | 7.4858 |
| 3_MDA-MB-436<sup>7TGP-POS</sup> Upregulated | Low | 11.2013 | 17.8621 | 11.832 | 11.2013 |
| 4_MDA-MB-436<sup>7TGP-POS</sup> Downregulated | Low | 6.1634 | 9.3091 | 7.0964 | 6.1634 |
| 5_MDA-MB-231<sup>7TGP-POS</sup> Upregulated | Standard | 9.9889 | 13.3504 | 13.5261 | 9.9889 |
| 6_MDA-MB-231<sup>7TGP-POS</sup> Downregulated | Standard | 8.1561 | 14.7582 | 5.48 | 5.48 |
| 7_MDA-MB-436<sup>7TGP-POS</sup> Upregulated | Standard | 10.6394 | 17.4561 | 11.3186 | 10.6394 |
| 8_MDA-MB-436<sup>7TGP-POS</sup> Downregulated | Standard | 6.0908 | 8.0855 | 10.1611 | 6.0908 |

For each gene set, the smallest HSD statistic value across MLIA vs. LAR, BLIA, and BLIS comparisons was used to summarize its ability to distinguish MLIA from other subtypes. Overall, gene sets composed of upregulated genes exhibited higher HSD statistic minimum values than their downregulated counterparts, indicating that Wnt-activated transcriptional programs more robustly distinguish MLIA tumors. For example, the upregulated gene sets from MDA-MB-231^7TGP-POS^ and MDA-MB-436^7TGP-POS^ cells yielded HSD minimum values ranging from 9.29 to 11.20, whereas corresponding downregulated gene sets ranged from 5.48 to 7.49. Notably, the upregulated gene set from MDA-MB-436^7TGP-POS^ cells cultured in low serum exhibited the highest minimum HSD value (11.20), indicating the strongest separation of MLIA from other subtypes. Although upregulated gene sets generally exhibited stronger subtype discrimination, an exception was observed for MDA-MB-231^7TGP-POS^ cells, where downregulated genes produced higher HSD values than upregulated ones in the MLIA vs. BLIA comparison. This suggests that Wnt-repressed genes may also contribute to subtype-specific transcriptional differences in certain contexts. Low serum-derived gene sets generally exhibited equal or higher HSD values compared to their standard serum counterparts, particularly in the MDA-MB-436 models, suggesting that serum deprivation may accentuate Wnt-driven transcriptional differences aligned with TNBC subtype variation.

Our findings demonstrate that Wnt-related transcriptional programs derived from TNBC cell lines are not only preserved in patient tumors but are particularly enriched in the MLIA subtype, highlighting a robust association between Wnt activation and this TNBC variant. The ability of these gene sets to stratify TNBC subtypes while capturing key pro-tumorigenic processes underscores their potential for molecular classification and therapeutic targeting in Wnt-responsive TNBC.

## DISCUSSION

Aberrant activation of the Wnt signaling pathway in TNBC has been implicated in drug resistance, recurrence, and metastasis. In this bioinformatics study, we investigated the transcriptional consequences of endogenous Wnt activity in TNBC cells and its relationship to clinically relevant subtypes. We first measured basal Wnt activity (in the absence of exogenous Wnt ligands) using the 7TGP Tcf/Lef reporter vector, and subsequently generated Wnt activity reporter TNBC cell lines. These cellular models were then used to profile transcriptomic changes under low and standard nutrient conditions. By leveraging these paired cell lines, we identified Wnt-associated transcriptional signatures that operate under basal conditions across distinct nutrient environments.

Previously, a study using the 7TGP vector demonstrated that MDA-MB-231 cells, when overexpressing E-cadherin, exhibited higher Wnt activity (eGFP levels) compared to parental MDA-MB-231 cells^36^. Another study derived the Wnt activity reporter TNBC cell line MDA-MB-468-TOPGFP using the 7TGP vector and reported that treatment with CHIR99201 increased Wnt activity (eGFP levels) and reduced sensitivity to carboplatin^37^. Therefore, the Wnt activity cell lines generated and validated in our study could be employed to evaluate how Wnt activation could influence platinum-based compounds or other chemotherapies administered to TNBC patients, potentially contributing to drug resistance. Although identifying potent Wnt inhibitors has been challenging^38^, these cell lines may serve as valuable tools for discovering novel Wnt pathway inhibitors through compound library screening.

An important factor we considered when evaluating Wnt activity was the serum concentration in the culture media. We aimed to compare the transcriptomic profiles of Wnt-positive and Wnt-negative TNBC cell lines under nutrient-poor and nutrient-rich conditions. Serum deprivation mimics aspects of the tumor microenvironment, such as hypovascularized regions where growth factors are limited, thereby amplifying pathway-specific transcriptional responses. In a previous study, our group identified Wnt target genes in TNBC cells following Wnt3A stimulation^25^. In the current study, the 7TGP reporter enabled us to detect basal Wnt activity under both low and standard serum conditions, without requiring exogenous ligand stimulation. Our findings suggest that serum availability modulates Wnt-driven transcriptional outputs, highlighting the importance of physiological context for in vitro models of TNBC. Interestingly, metabolic stress induced by nutrient deprivation has been shown to modulate Wnt pathway activity and promote cancer cell plasticity in MDA-MB-231 cells^39^. Understanding how nutrient availability shapes Wnt signaling in TNBC cells may help optimize in vitro assays and better model Wnt-driven phenotypes relevant to the tumor microenvironment.

In addition to nutrient effects, we observed significant variability in Wnt activity both between and within cell lines. Even in clonal cell populations, a spectrum of Wnt activity exists, emphasizing the plasticity of TNBC cells and the importance of single-cell resolution approaches. The ability to isolate Wnt-positive and Wnt-negative subpopulations provided a unique opportunity to define Wnt-driven transcriptional programs under physiologically relevant serum conditions.

Gene enrichment analysis revealed that Wnt-positive TNBC cells upregulate pro-tumorigenic transcriptional programs, including cytokine responses, immune cell recruitment, EMT, and receptor tyrosine kinase signaling. Wnt-positive TNBC cells consistently exhibited enrichment of IL6/JAK/STAT3 signaling, TNFα signaling via NF- κB, and EMT hallmark gene sets under both serum conditions. These findings suggest that Wnt activity intrinsically links inflammatory responses with EMT and stemness in TNBC. Interestingly, IL-6 inhibition using Tocilizumab reduced the mesenchymal phenotype and stemness properties of MDA-MB-231 cells by inhibiting Wnt signaling and enhancing their sensitivity to cisplatin^40^.

GSEA also revealed upregulation of cholesterol homeostasis pathways in Wnt-negative TNBC cells, suggesting an inverse relationship with Wnt activity^41^. Furthermore, the Notch signaling hallmark was enriched in MDA-MB-436^7TGP-POS^ cells, while MTORC1 signaling was activated in MDA-MB-436^7TGP-NEG^ cells. These findings highlight potential crosstalk between Wnt, Notch, and mTORC1 signaling axes, with implications for combination therapy strategies for TNBC.

Furthermore, a 55-gene expression Wnt signature comprising 42 upregulated and 13 downregulated genes, captured core aspects of Wnt activation across both Wnt-positive cell lines and both serum conditions. This signature captured transcriptional programs associated with EMT, inflammation, GPCR signaling, and ECM remodeling. Network and protein–protein interaction analysis further identified chemokines, receptors, and ECM components as central hubs. PPI analysis findings linking *CXCL1*, *CXCL8*, and *CX3CL1* expression with Wnt-positive TNBC cells suggest that Wnt signaling may drive inflammatory chemokine programs that promote EMT and stemness, reinforcing aggressive phenotypes and potentially shaping an immune-altered tumor microenvironment. This notion is supported by studies showing that breast cancer stem cell properties are enhanced by *CXCL1*^42,43^ as well as *CXCL8*^44,45^ expressions.

When the Wnt-derived gene sets were assessed in a large TNBC RNA-Seq cohort (n = 699), MLIA tumors consistently exhibited the highest expression scores across nearly all signatures and derived clusters. The cytokine signaling enrichment observed in Wnt-positive cells may reflect the immune-altered phenotype of MLIA tumors. These parallels underscore the Wnt pathway’s role in shaping the mesenchymal identity of TNBC and suggest that targeting Wnt activity could complement existing therapeutic strategies aimed at mitigating stem-like tumor traits. Our findings support a model in which Wnt signaling contributes to the transcriptional identity of MLIA tumors and may serve as a functional marker of this molecular TNBC subtype.

Together, our findings define Wnt-responsive gene sets that stratify TNBC subtypes and reveal candidate biomarkers and therapeutic targets for Wnt-directed interventions. By integrating cell line–derived transcriptional programs with patient tumor datasets, we demonstrate that Wnt activity is particularly enriched in the MLIA molecular subtype of TNBC and associated with pro-tumorigenic features such as EMT, inflammation, and stemness. These insights lay the groundwork for therapeutic strategies aimed at modulating Wnt signaling and raise the possibility that Wnt pathway activity may influence tumor–immune interactions relevant to immunotherapy responsiveness. Ultimately, targeting Wnt-driven processes could improve treatment efficacy in TNBC.

## Methods

### Cell Culture

The MDA-MB-231 cell line was a kind gift of Dr. Mina Bissell (University of California, Berkeley, CA, USA) and the MDA-MB-436 cell line was purchased from American Type Culture Collection (ATCC, LGC Promochem, Molsheim, France). Both cell lines were authenticated in 2021 by short tandem repeat profiling using the Powerplex 16 system (Promega, Charbonnières-les-Bains, France). MDA-MB-231 cells are mutated for *TP53*, *KRAS*, and *CDKN2A*, while MDA-MB-436 cells are mutated for *TP53* and *BRCA1, and* deficient for PTEN^6,46,47^. MDA-MB-231, MDA-MB-231^7TGP^, MDA-MB-231^7TGP-NEG^, and MDA-MB-231^7TGP-POS^ cells were maintained in complete DMEM/F-12 media (Life Technologies, Courtaboeuf, France) supplemented with 10% (vol/vol) fetal bovine serum (FBS, Life Technologies), 100 U/mL penicillin, and 100 µg/mL streptomycin (P/S, Life Technologies). MDA-MB-436, MDA-MB-436^7TGP^, MDA-MB-436^7TGP-NEG^, and MDA-MB-436^7TGP-POS^ cells were cultured in complete RPMI-1640 (Life Technologies) supplemented with 10% (vol/vol) FBS (Life Technologies), 100 U/mL penicillin, and 100 µg/mL streptomycin (P/S, Life Technologies).

Briefly, tumor cells were passaged by first collecting the culture supernatants, rinsing the culture flask with 1X PBS (no calcium, no magnesium), adding the 1X PBS-cell suspension to the collected culture media, and incubating the culture flasks with a corresponding amount of TrypLE Express (LifeTechnologies) for 2-3 minutes; once the cells detached, the flasks were rinsed with 1X PBS (no calcium, no magnesium) and the cell-containing TryPLE Express/PBS suspension was added to the previous supernatant collection; the resultant cell suspension was centrifuged at 1500 RPM for 5 minutes, the supernatant was discarded, and the pellet of harvested cells was resuspended in fresh culture media to be expanded in new flasks (referred from herein as TryPLE Express harvesting protocol). Cells were maintained at 37.5°C, with humidity, and 5% carbon dioxide.

### Generation of MDA-MB-231^7TGP^ and MDA-MB-436^7TGP^ Tcf/Lef reporter cell lines

The 7TGP vector was a kind gift from Roel Nusse (Addgene plasmid # 24305). To expand the 7TGP vector, One Shot Stbl3 *E. coli* cells (Life Technologies) were transformed with the 7TGP vector according to the manufacturer’s instructions, and the 7TGP plasmid was isolated from transformed Stbl3 *E. coli* cells using the NucleoSpin Plasmid Mini kit for plasmid isolation (Macherey-Nagel, Hœrdt, France). Desalted plasmid stocks (>1 µg / µl) were sequenced to verify DNA sequence integrity. The desalted 7TGP vector was then transfected into MDA-MB-231 or MDA-MB-436 cells using the MDA-MB- 231 Cell Avalanche Transfection Reagent (EZ Biosystems, College Park, MD, USA) following the manufacturer’s instructions. Puromycin selection (3 µg / mL) was initiated 48 hours after transfection and cells were kept under puromycin selection (3 µg / mL) for five passages. The resultant cell lines, MDA-MB-231^7TGP^ and MDA-MB-436^7TGP^, were deemed as stably transfected with the 7TGP vector.

### Flow cytometry analyses and FACS

A ZE5 Cell Analyzer (Bio-Rad, Hercules, CA, USA) was used to assess the frequency and Mean Fluorescent Intensity (MFI) of eGFP from the 7TGP reporter in the MDA-MB-231^7TGP^ and MDA-MB-436^7TGP^ cell lines, as well as to assay cell viability using a fixable viability dye (live unstained, dead stained), either Zombie Violet or Zombie NIR (Biolegend, San Diego, CA, USA). A minimum of 30,000 total events were collected for flow cytometric analyses. An S3 fluorescence-activated cell sorter (Bio-Rad) was used to enrich eGFP-negative and eGFP-positive cells to establish the following cell lines: MDA- MB-231^7TGP-NEG^, MDA-MB-231^7TGP-POS^, MDA-MB-436^7TGP-NEG^, and MDA-MB-436^7TGP-POS^ cells. The sorting strategy involved culturing MDA-MB-231^7TGP^ and MDA-MB-436^7TGP^ cells in 10% FBS complete media, isolating the top third of eGFP-positive cells (FL1 channel) and the bottom third of eGFP-negative cells (leftmost on FL1), expanding each population, and repeating this sort-and-expand cycle seven additional times.

### Cellular assays

Frequency and MFI of the eGFP from the 7TGP reporter in the MDA-MB-231^7TGP^ and MDA-MB-436^7TGP^ were first measured under standard conditions (10% FBS complete media, 37.5°C, with humidity, and 5% carbon dioxide) using the ZE5 Cell Analyzer (Bio-Rad). To test the effects of Wnt agonist CHIR99021 (Selleckchem) on the levels of eGFP in the MDA-MB-231^7TGP^ and MDA-MB-436^7TGP^ cells, cells were harvested from standard condition cultures using the TrypleExpres harvest protocol, and culture media serum was washed off twice with 1X PBS (no calcium or magnesium and changing each time the conical polypropylene tube); serum-free cells were then seeded at 70% confluence in 0.5% FBS complete media and incubated for 48 hours in this minimal nutrient condition, after which, the Wnt agonist CHIR99021 or DMSO was added and incubated for 24 hours before being harvested using the TryPLE Express harvest protocol for analyses using the ZE5 Cell Analyzer (Bio-Rad). The agonist CHIR99021 (dissolved in DMSO) was used at a concentration of 3 µM and compared to an equal volume of DMSO as a control.

### RNA-Seq and bioinformatic analyses

The established MDA-MB-231^7TGP-NEG^, MDA-MB-231^7TGP-POS^, MDA-MB-436^7TGP-NEG^, and MDA-MB-436^7TGP-POS^ cell lines were harvested from 10% FBS complete media cultures using the TryPLE Express harvest protocol and then cultured in low (0.5% FBS) or standard (10% FBS) serum complete media for subsequent RNA-Seq analyses.

For low serum cultures, harvested cells were rinsed twice with 1X PBS (no calcium or magnesium and changing each time the conical polypropylene tube), and cells were then seeded in T75 culture flasks at 70% confluency in low serum media (0.5% FBS). On day 3 post-seeding, cells were harvested by collecting the supernatant, gently scraping the cell layer in 1X PBS (no calcium, no magnesium), adding the PBS-cell suspension to the collected supernatant, centrifuging the resultant cell suspension at 1500 RPM for 5 minutes, and aspirating the supernatant to freeze the cell pellets.

For standard serum cultures, harvested cells were seeded at low density in T75 culture flasks using standard serum media (10% FBS) until reaching 70% confluency by day 6 post-seeding. Cells were then harvested by collecting the supernatant, gently scraping the cell layer in 1X PBS (no calcium, no magnesium), adding the PBS-cell suspension to the collected supernatant, centrifuging the resultant cell suspension at 1500 RPM for 5 minutes, and aspirating the supernatant and freezing the cell pellets.

Cell pellets were stored at -80°C until RNA extraction was performed using the RNAeasy Mini Kit (Qiagen, Courtaboeuf, France), according to the manufacturer’s instructions. RNA quantity and quality (260/280, 260/230 ratios) were measured using a NanoDrop UV-Vis spectrophotometer (Thermo Fisher Scientific, France) and verified using a Bioanalyzer (Agilent Technologies). All samples were subjected to quality control on a Bioanalyzer instrument and most total RNA used for sequencing showed a RIN (RNA Integration Number) > 7. RNA-Seq analyses for each experimental condition were done on biological duplicates.

Total RNA was extracted from low serum cultures and sent to MedGenome (Bangalore, India) for RNA sequencing. Libraries were prepared using 1 µg of RNA and the NEBNext® Ultra™ II Directional RNA Library Prep Kit for Illumina (NEB #E7775). Ribosomal RNA was removed from 500 ng of total RNA using biotinylated, target-specific oligos and magnetic beads (RiboCop rRNA Depletion Kit, Lexogen). The rRNA-depleted RNA was fragmented using divalent cations under elevated temperature and reverse transcribed into first-strand cDNA using random hexamer primers and reverse transcriptase. Second-strand synthesis was performed using DNA Polymerase I and RNase H. End repair, A-tailing, and adaptor ligation were then performed. Adapter-ligated fragments were PCR-amplified under the following conditions: initial denaturation at 98°C for 30 s; 12 cycles of 98°C for 10 s and 65°C for 75 s; and final extension at 65°C for 5 min. Final libraries were purified, analyzed on the Fragment Analyzer using the High Sensitivity NGS Fragment Kit (Agilent, DNF-474-1000), and quantified using the Qubit dsDNA HS Assay Kit (Invitrogen, Q32854). Libraries were pooled, diluted to optimal loading concentrations, and sequenced on the Illumina HiSeq X system using 2 × 150 bp paired-end reads, generating ∼80 million clusters per sample.

From standard serum cultures, RNA sequencing libraries were prepared from 1 µg of total RNA using the Illumina TruSeq Stranded mRNA Library preparation kit which enables strand-specific RNA sequencing. A first step of polyA selection using magnetic beads was done to sequence polyadenylated transcripts. After fragmentation, cDNA synthesis was performed, and resulting fragments were used for dA-tailing and then ligated to the TruSeq indexed adapters. PCR amplification was finally achieved to create the final cDNA library. Individual library quantification and quality assessment were performed using Qubit fluorometric assay (Invitrogen) with dsDNA HS (High Sensitivity) Assay Kit and LabChip GX Touch using a High Sensitivity DNA chip (Perkin Elmer). Libraries were then equimolarly pooled and quantified by qPCR using the KAPA library quantification kit (Roche). Sequencing was carried out on the NovaSeq 6000 instrument from Illumina using paired-end 2 x 100 bp, to obtain around 50 million clusters per sample. The Q30 value for each RNA sample sequenced was above 80%.

The overall quality of the reads was first checked using FastQC (v0.11.8) and the sequencing orientation, also known as strandness, was assessed using RseQC (v2.6.4). Those reads were then aligned on both a ribosomal RNAs database using Bowtie (v1.2) and on the human reference genome (hg19 assembly) using STAR (v2.6.1b). Raw counts were generated using STAR gene quantification mode on the Gencode v19 gene database. Coding genes with almost null expression (less than 10 reads among all samples) were filtered out. Filtered counts were then normalized using the R package DESeq2 (v1.26.0) ^48^. We were able to detect the expression of a total of 15,512 genes for the low serum samples and 14,899 genes for the standards serum samples. Variance-stabilizing transformation was then applied to perform exploratory analysis using the R package FactoMineR (v1.42) ^49^ and heat map visualizations were created using the R package pheatmap (v1.0.12). Differential expression files were also generated with the R package DESeq2 (v1.26.0) using an adjusted P value of 5% and an absolute log2 fold change of 1. Gene Ontology (GO) enrichment analysis was performed on deregulated genes (up- and down-deregulated mixed) with an absolute log2 fold change of 1 for biological process (BP) and molecular function (MF) ontologies using the R package clusterProfiler (v3.14.0). For GO BP analyses, bubble plots, bar graphs, and Venn diagrams were generated in Python (Matplotlib v3.8). In addition, GO MF and pathway analyses were performed using Metascape ^50^. KEGG pathway analysis ^51^ was also performed on the deregulated genes using ShinyGO 0.82 (http://bioinformatics.sdstate.edu/go/) ^52^ and the Wnt signaling pathway schematics were rendered using Pathview ^53^. Gene Set Enrichment Analysis (GSEA) software ^54,55^ and the Hallmark gene sets from Molecular Signature Database (MSigDB) ^56,57^ were used to analyze all the genes (15,512 genes for low serum conditions and 14,899 genes for standard serum conditions) that were pre-ranked by log2 fold change, as it is recommended to perform GSEA using unfiltered gene lists.

### TNBC cohort analyses

The enrichment of the gene expression signatures derived from the Wnt-negative and Wnt-positive TNBC cell lines was tested in a RNA-Seq TNBC cohort (n = 699), which is comprised of four molecular subtypes (Luminal Androgen Receptor, Mesenchymal-like Immune-Altered, Basal-like Immune-Activated, Basal-like Immune-Suppressed)^9^. Gene expression scores were calculated using the formula:

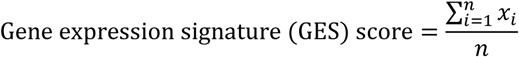

where *i* represents a gene indicator, *x* is the expression value of each gene, and *n* is the total number of genes in the signature. This formula averages the expression values of all genes within a given signature to determine the GES score for each sample.

Heatmap visualizations were generated to depict the expression patterns of Wnt-related gene signatures across the four TNBC molecular subtypes and determine differential gene clusters amongst the four molecular subtypes of the RNA-Seq TNBC cohort. Differential gene clusters were analyzed using gene ontology enrichment analyses (GOEA)^58^ for GO BP.

### Statistical analyses

For the cellular assays, the significance of the results was tested in GraphPad Prism (GraphPad) using an unpaired two-tailed Student’s *t*-test; significance was represented as * *P* < 0.05, ** *P* < 0.01, *** *P* < 0.001, and error bars represent standard deviation. For the gene enrichment signature score analyses, statistical analysis was performed using one-way analysis of variance (ANOVA) with Tukey post hoc test; P values ≤ 0.01 were considered significant. Stouffer’s method was used to combine P values across low and standard serum conditions, allowing identification of the top GO BP categories consistently enriched in both. In order to identify differential gene clusters, hierarchical clustering with centered Pearson’s correlation and Ward’s linkage method, and with Euclidean distance and Ward’s linkage method, was performed for genes and samples, respectively. Pearson correlation analyses were conducted to evaluate associations between Wnt-related gene expression signatures and lists of differential gene clusters. Tukey’s Honestly Significant Difference (HSD) statistic was used to identify significant differences in expression of gene expression signatures derived from Wnt-negative and Wnt-positive TNBC cell lines across the four molecular TNBC subtypes.

### Availability of data

The RNA-Seq datasets have been deposited in Gene Expression Omnibus and will be available once the manuscript will be accepted for publication. Gene Ontology data are provided in the supplementary files.

## Acknowledgments

RNA-Seq data management, quality control, and primary analysis were performed by the Bioinformatics platform of the Institut Curie. We are grateful to Dr Samyuktha Suresh’s critical reading of the manuscript. We acknowledge Biorender for the creation of schematics.

## Author Contribution Statement

**Ramón García-Areas:** Conceptualization, Methodology, Software, Validation, Formal analysis, Investigation, Data curation, Writing—original draft preparation, Writing—review and editing, Visualization. **Elodie Girard:** Software, Validation, Formal analysis, Data curation, Writing—review and editing, Visualization. **Hamza Lasla:** Software, Validation, Formal analysis, Data curation, Writing—review and editing, Visualization. **Virginie Raynal:** Formal analysis, Data curation, Writing—review and editing. **Amit Kumar Pandey:** Formal analysis, Data curation, Writing—review and editing. **Nicolas Servant:** Formal analysis, Data curation, Writing—review and editing, Supervision. **Pascal Jézéquel:** Formal analysis, Data curation, Writing—review and editing, Supervision. **Thierry Dubois:** Conceptualization, Validation, Formal analysis, Data curation, Writing— original draft preparation, Writing—review and editing, Visualization, Supervision, Project administration, Funding acquisition.

## Appendix A. Supplementary Files

Supplementary File 1: Complete lists of genes differentially expressed between Wnt-positive and Wnt-negative TNBC cell lines;

Supplementary File 2: Lists of genes previously reported as Wnt-related

Supplementary File 3: Complete lists and overlap of GO BP Terms enriched by genes deregulated in Wnt-positive and Wnt-negative TNBC cell lines;

Supplementary File 4: Complete lists of GO MF Terms enriched by genes deregulated in Wnt-positive and Wnt-negative TNBC cell lines;

Supplementary File 5: KEGG analyses of genes differentially expressed by Wnt-negative and Wnt-positive TNBC cell lines;

Supplementary File 6: Unfiltered lists of genes expressed by Wnt-negative and Wnt-positive TNBC cell lines;

Supplementary File 7: Hallmark MSigDB GSEA analyses on genes expressed by Wnt-negative and Wnt-positive TNBC cell lines.

Supplementary File 8: Data Summary for TNBC Patient Data Analyses

Supplemental File 9: Gene list for each GES including H1/H2 gene lists and H1/H2 GO BP analyses

