## Supplementary Figures 1-11 for "Wnt signaling promotes inflammation and EMT-associated gene expression in mesenchymal TNBC"

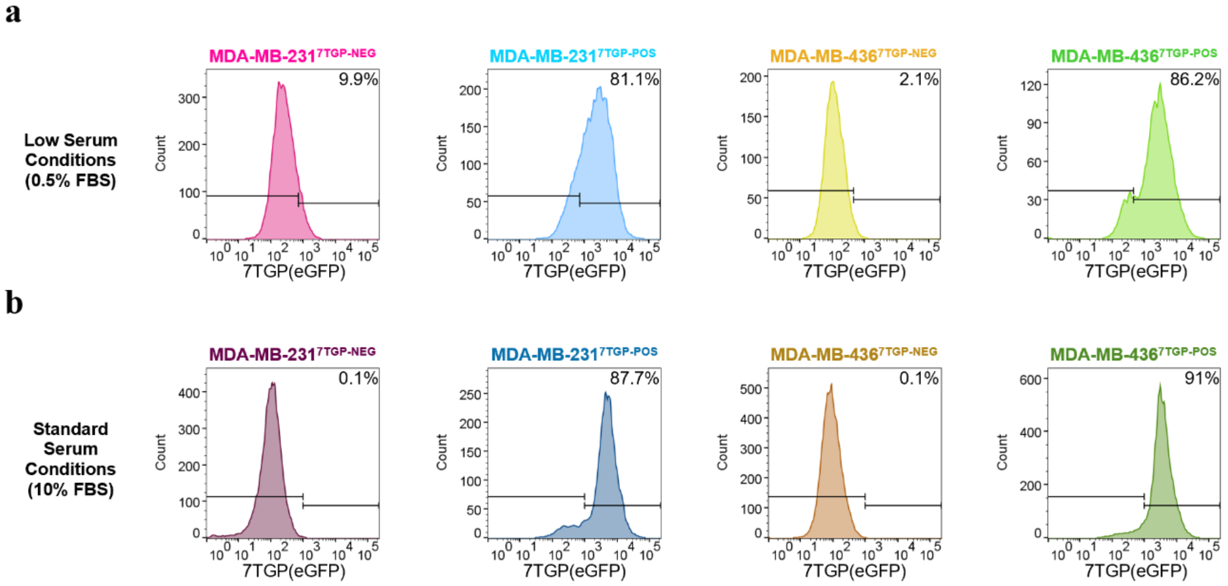


**Supplementary Fig. 1 | eGFP fluorescence profiles of Wnt-negative and Wnt-positive TNBC cell lines cultured in low or standard serum conditions. a, b** Representative flow cytometry histograms showing eGFP fluorescence in MDA-MB-231$^{7TGP-NEG}$, MDA-MB-231$^{7TGP-POS}$, MDA-MB-436$^{7TGP-NEG}$, and MDA-MB-436$^{7TGP-POS}$ cell lines after culture for 3 days in low serum (a) or 6 days in standard serum (b).

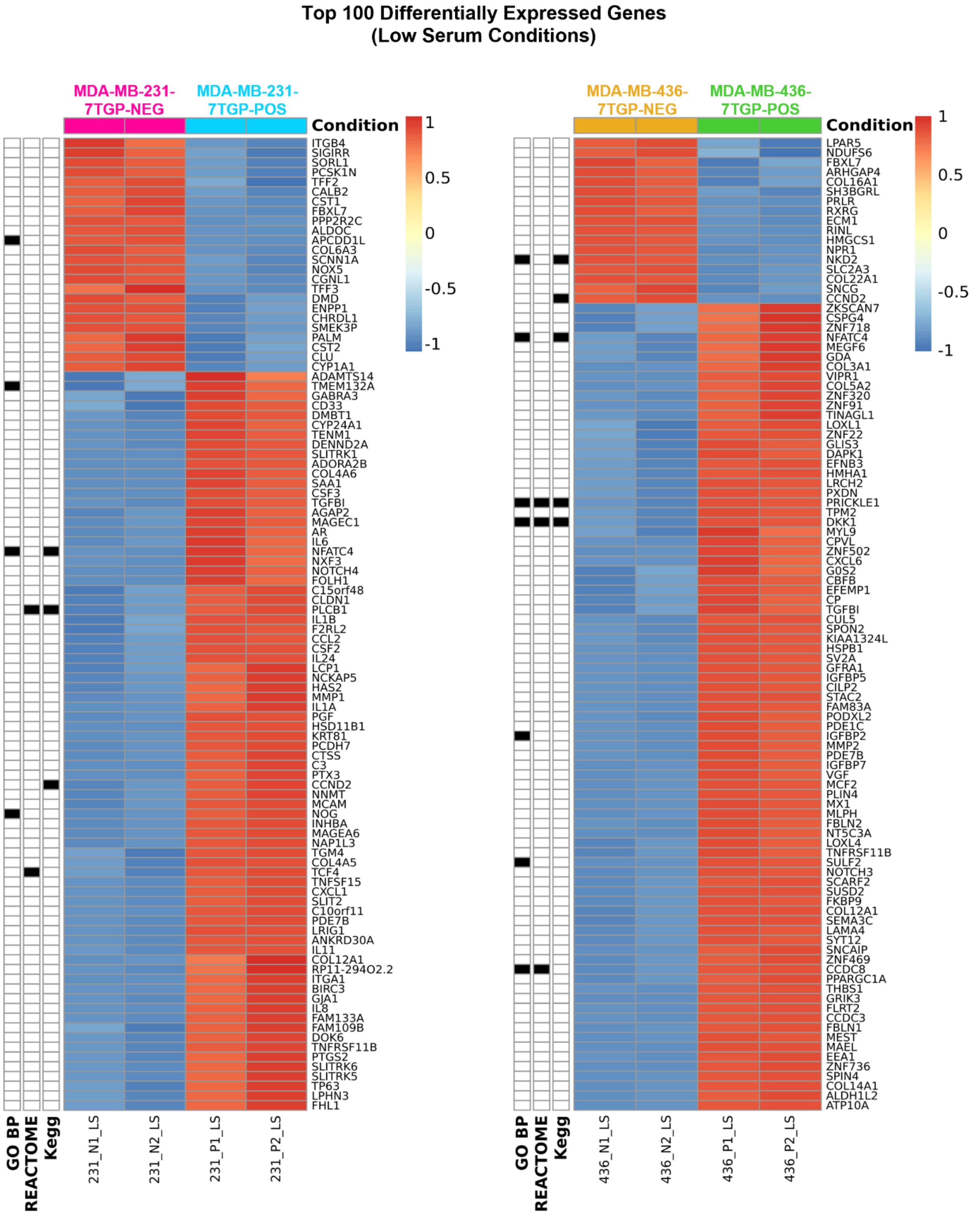
Top 100 Differentially Expressed Genes
(Low Serum Conditions)
MDA-MB-231-7TGP-NEG
MDA-MB-231-7TGP-POS
Condition
1
0.5
0
-0.5
-1
ITGB4
SIGIRR
SORL1
PCSK1N
TFF2
CALB2
CST1
FBXL7
PPP2R2C
ALDOC
APCDD1L
COL6A3
SCNN1A
NOX5
CGNL1
TFF3
DMD
ENPP1
CHRDL1
SMEK3P
PALM
CST2
CLU
CYP1A1
ADAMTS14
TMEM132A
GABRA3
CD33
DMBT1
CYP24A1
TENM1
DENND2A
SLITRK1
ADORA2B
COL4A6
SAA1
CSF3
TGFBI
AGAP2
MAGEC1
AR
IL6
NFATC4
NXF3
NOTCH4
FOLH1
C15orf48
CLDN1
PLCB1
IL1B
F2RL2
CCL2
CSF2
IL24
LCP1
NCKAP5
HAS2
MMP1
IL1A
PGF
HSD11B1
KRT81
PCDH7
CTSS
C3
PTX3
CCND2
NNMT
MCAM
NOG
INHBA
MAGEA6
NAP1L3
TGM4
COL4A5
TCF4
TNFSF15
CXCL1
SLIT2
C10orf11
PDE7B
LRIG1
ANKRD30A
IL11
COL12A1
RP11-294O2.2
ITGA1
BIRC3
GJA1
IL8
FAM133A
FAM109B
DOK6
TNFRSF11B
PTGS2
SLITRK6
SLITRK5
TP63
LPHN3
FHL1
GO BP
REACTOME
Kegg
231_N1_LS
231_N2_LS
231_P1_LS
231_P2_LS
MDA-MB-436-7TGP-NEG
MDA-MB-436-7TGP-POS
Condition
1
0.5
0
-0.5
-1
LPAR5
NDUFS6
FBXL7
ARHGAP4
COL16A1
SH3BGRL
PRLR
RXRG
ECM1
RINL
HMGCS1
NPR1
NKD2
SLC2A3
COL22A1
SNCG
CCND2
ZKSCAN7
CSPG4
ZNF718
NFATC4
MEGF6
GDA
COL3A1
VIPR1
COL5A2
ZNF320
ZNF91
TINAGL1
LOXL1
ZNF22
GLIS3
DAPK1
EFNB3
HMHA1
LRCH2
PXDN
PRICKLE1
TPM2
DKK1
MYL9
CPVL
ZNF502
CXCL6
G0S2
CBFB
EFEMP1
CP
TGFBI
CUL5
SPON2
KIAA1324L
HSPB1
SV2A
GFRA1
IGFBP5
CILP2
STAC2
FAM83A
PODXL2
PDE1C
IGFBP2
MMP2
PDE7B
IGFBP7
VGF
MCF2
PLIN4
MX1
MLPH
FBLN2
NT5C3A
LOXL4
TNFRSF11B
SULF2
NOTCH3
SCARF2
SUSD2
FKBP9
COL12A1
SEMA3C
LAMA4
SYT12
SNCAIP
ZNF469
CCDC8
PPARGC1A
THBS1
GRIK3
FLRT2
CCDC3
FBLN1
MEST
MAEL
EEA1
ZNF736
SPIN4
COL14A1
ALDH1L2
ATP10A
GO BP
REACTOME
Kegg
436_N1_LS
436_N2_LS
436_P1_LS
436_P2_LS

**Supplementary Fig. 2 | Differential mRNA expression between Wnt-negative and Wnt-positive TNBC cell lines cultured under low serum conditions.**Heat maps generated from RNA-Seq data showing the top 100 DEGs between MDA-MB-231$^{7TGP\text{-}POS}$ and MDA-MB-231$^{7TGP\text{-}NEG}$ cells (left panel) and between MDA-MB-436$^{7TGP\text{-}POS}$ and MDA-MB-436$^{7TGP\text{-}NEG}$ cells (right panel), following culture for 3 days in low serum conditions (0.5% FBS). Genes previously identified as Wnt-related, based on three gene sets (GO Biological Process: cell–cell signaling by Wnt; Reactome: signaling by Wnt; KEGG: Wnt signaling pathway), are marked in black along the y-axis of each heat map. Each column represents a sample: MDA-MB-231 (231) or MDA-MB-436 (436); Wnt-negative (N) or Wnt-positive (P); replicate number (1, 2)

### Top 100 Differentially Expressed Genes
### (Standard Serum Conditions)

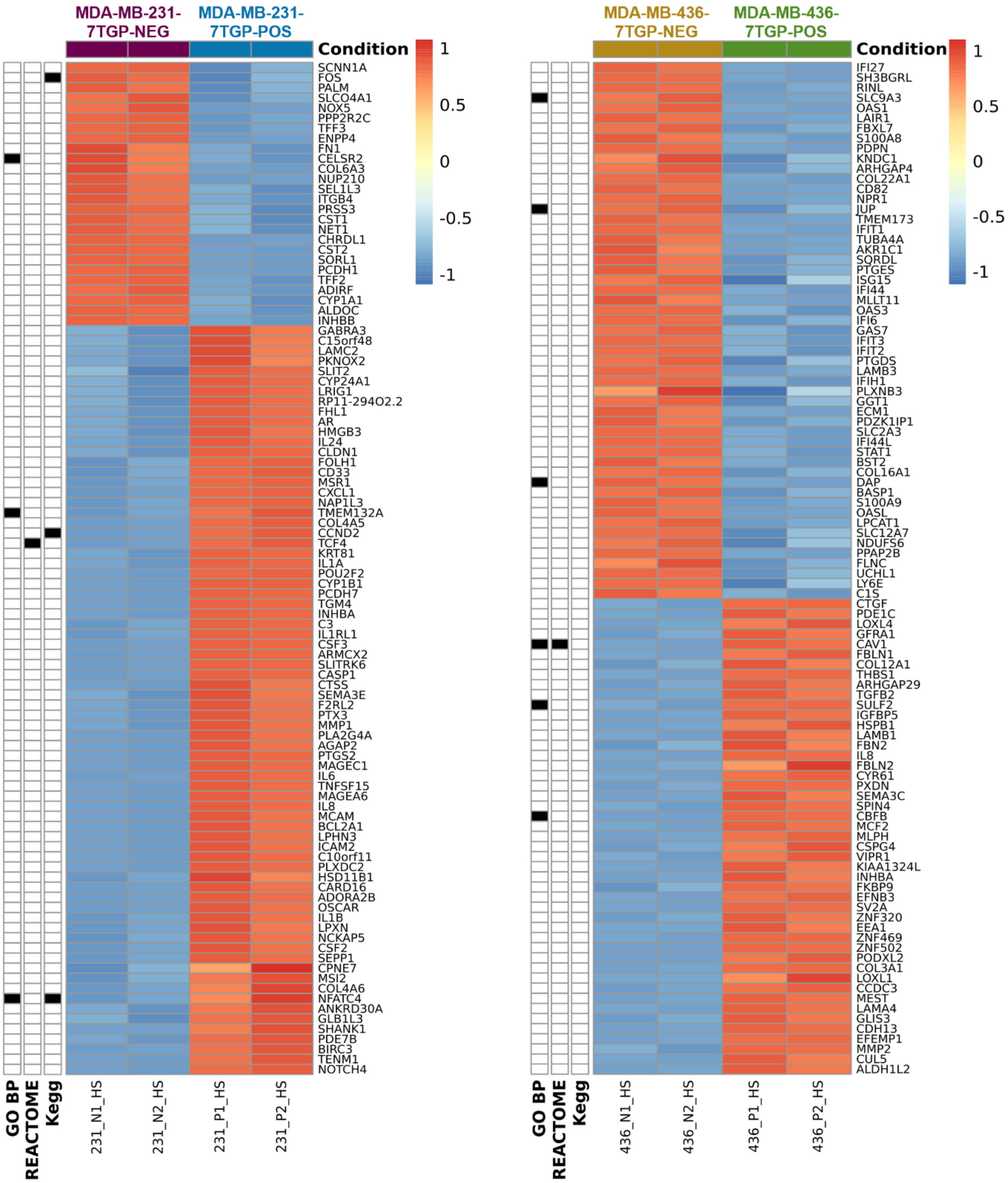

**Supplementary Fig. 3 | Differential mRNA expression between Wnt-negative and Wnt-positive TNBC cell lines cultured under standard serum conditions.** Heat maps generated from RNA-Seq data showing the top 100 DEGs between MDA-MB-231$^{7TGP\text{-}POS}$ and MDA-MB-231$^{7TGP\text{-}NEG}$ cells (left panel) and between MDA-MB-436$^{7TGP\text{-}POS}$ and MDA-MB-436$^{7TGP\text{-}NEG}$ cells (right panel), following culture for 3 days in standard serum conditions (10% FBS). Genes previously identified as Wnt-related, based on three gene sets (GO Biological Process: cell–cell signaling by Wnt; Reactome: signaling by Wnt; KEGG: Wnt signaling pathway), are marked in black along the y-axis of each heat map. Each column represents a sample: MDA-MB-231 (231) or MDA-MB-436 (436); Wnt-negative (N) or Wnt-positive (P); replicate number (1, 2).

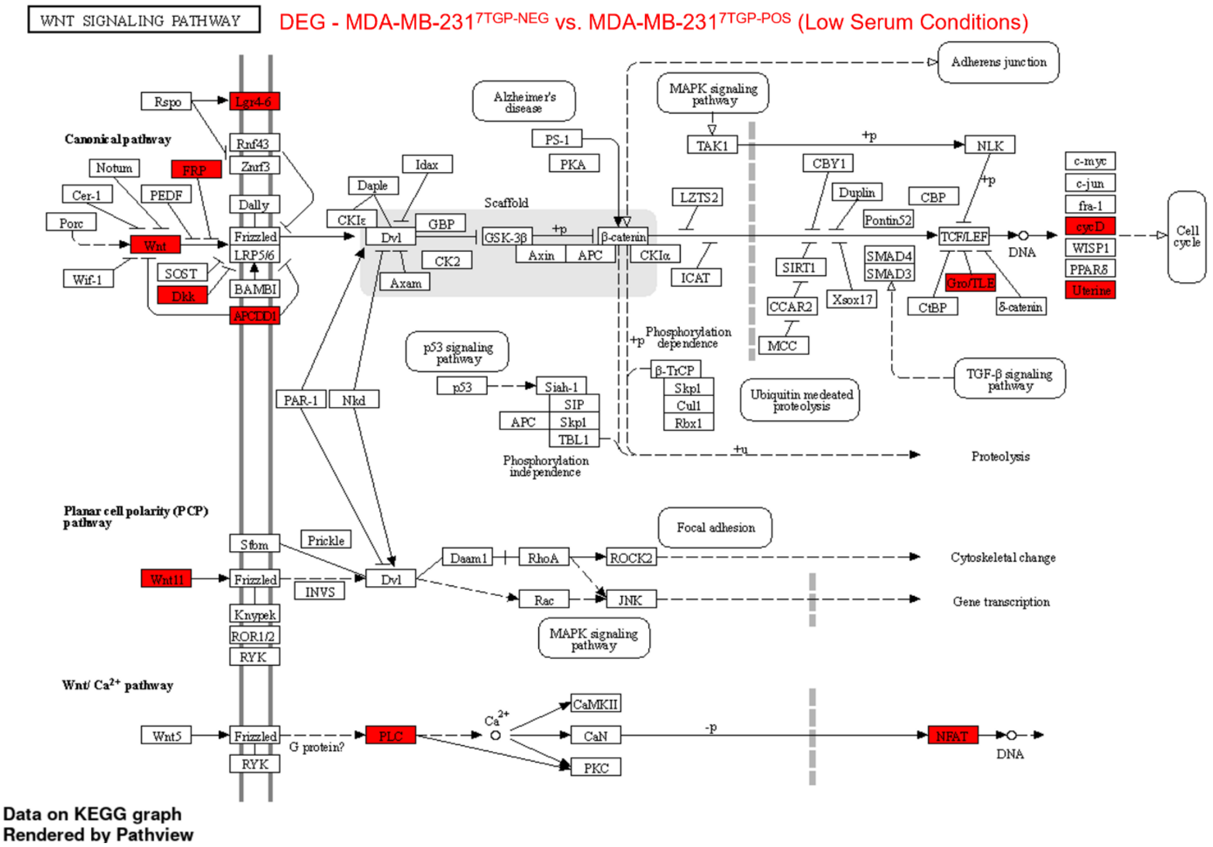


**Supplementary Fig. 4 | KEGG pathway map of Wnt signaling enriched by DEGs between MDA-MB-231$^{7TGP\text{-}NEG}$ and MDA-MB-231$^{7TGP\text{-}POS}$ cells.** The KEGG Wnt signaling pathway diagram highlights in red the genes or gene clusters differentially expressed between MDA-MB-231$^{7TGP\text{-}NEG}$ and MDA-MB-231$^{7TGP\text{-}POS}$ cells cultured in low serum. Visualization was generated using Pathview.

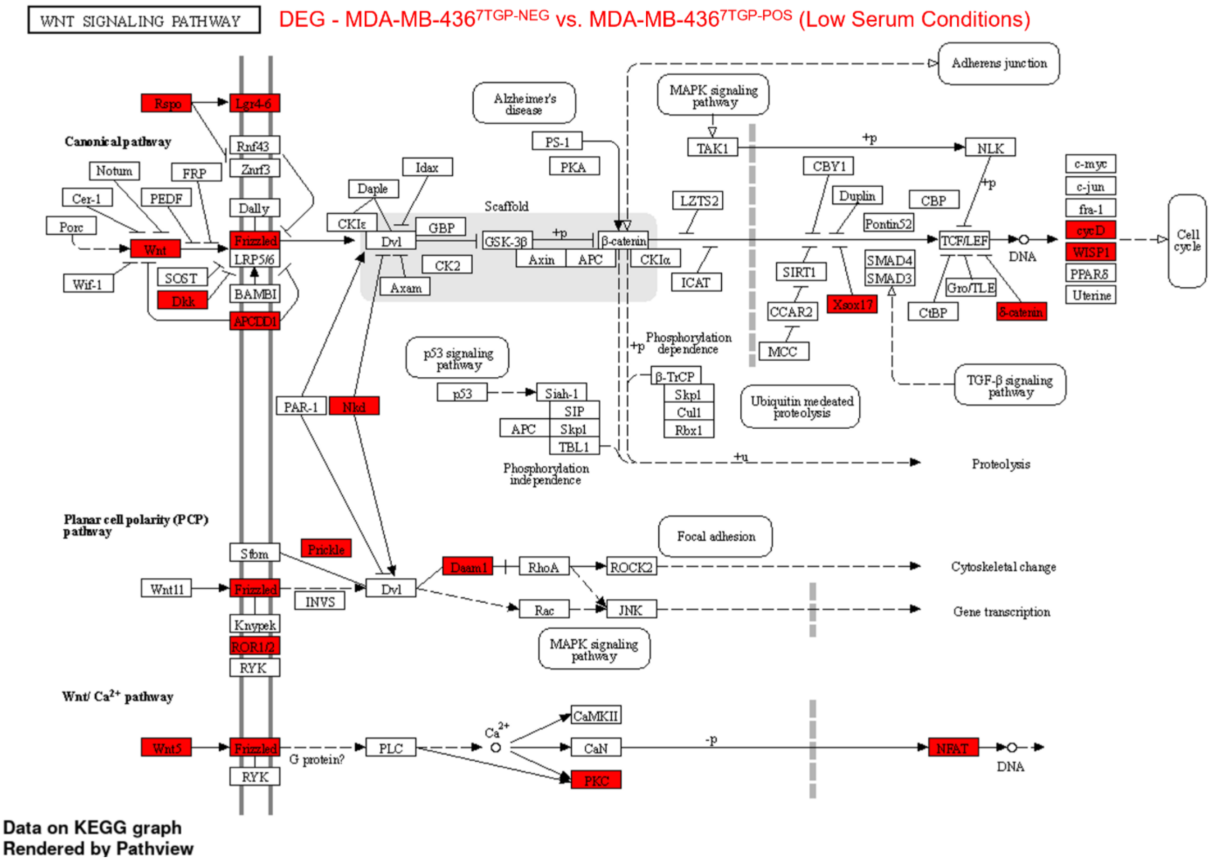


**Supplementary Fig. 5 | KEGG pathway map of Wnt signaling enriched by DEGs between MDA-MB-436$^{7TGP\text{-}NEG}$ and MDA-MB-436$^{7TGP\text{-}POS}$ cells.** The KEGG Wnt signaling pathway diagram highlights in red the genes or gene clusters differentially expressed between MDA-MB-436$^{7TGP\text{-}NEG}$ and MDA-MB-436$^{7TGP\text{-}POS}$ cells cultured in low serum. Visualization was generated using Pathview.

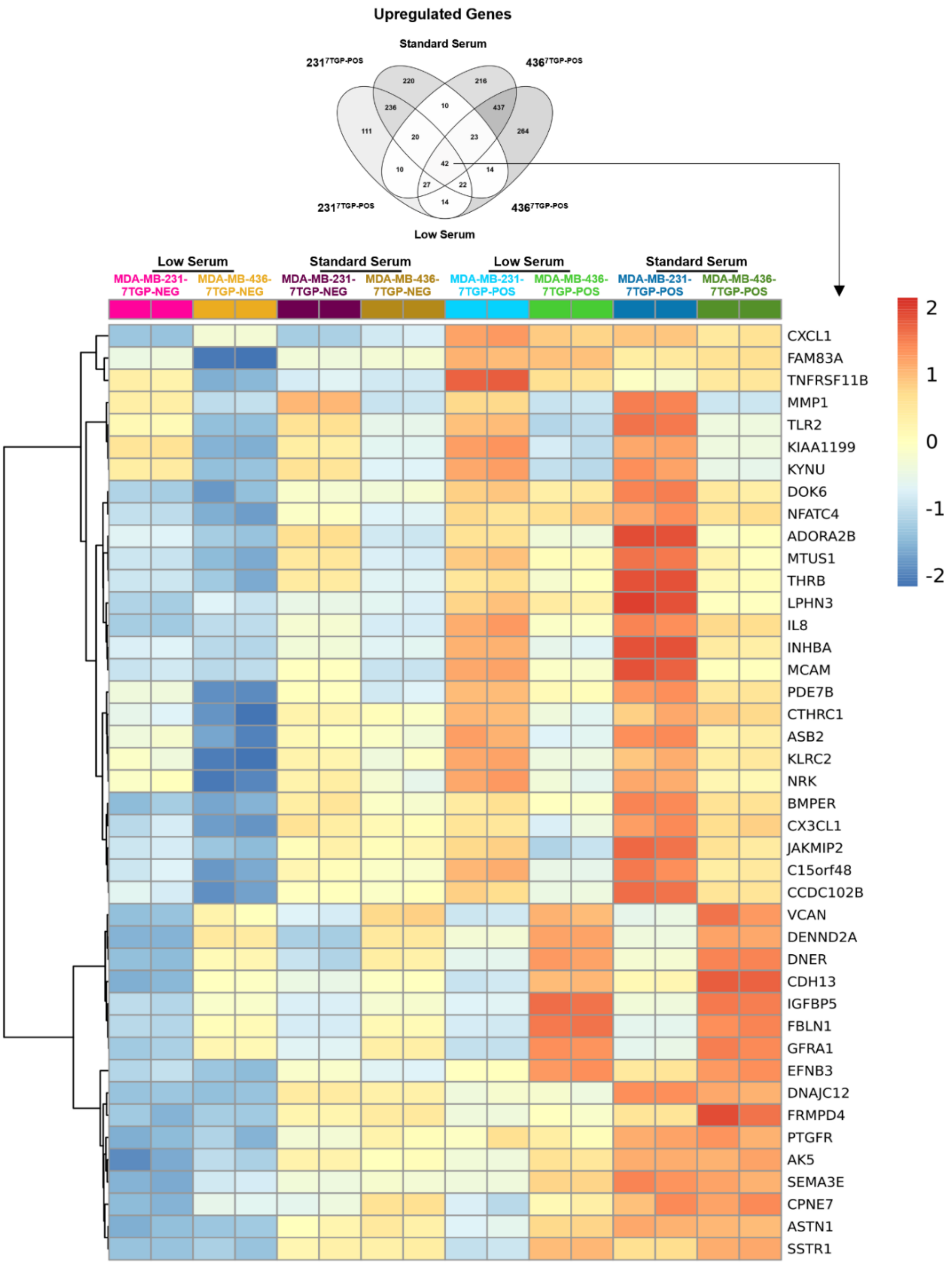


**Supplementary Fig. 6 | Shared upregulated genes across Wnt-positive TNBC cell lines and serum conditions.** Venn diagram showing the overlap of genes upregulated in Wnt-positive compared to Wnt-negative TNBC cells under both low and standard serum conditions. The heat map displays the expression patterns of the 42 genes consistently upregulated across all four comparisons. Differential expression was defined using an absolute $\log_2$ fold change ≥ 1

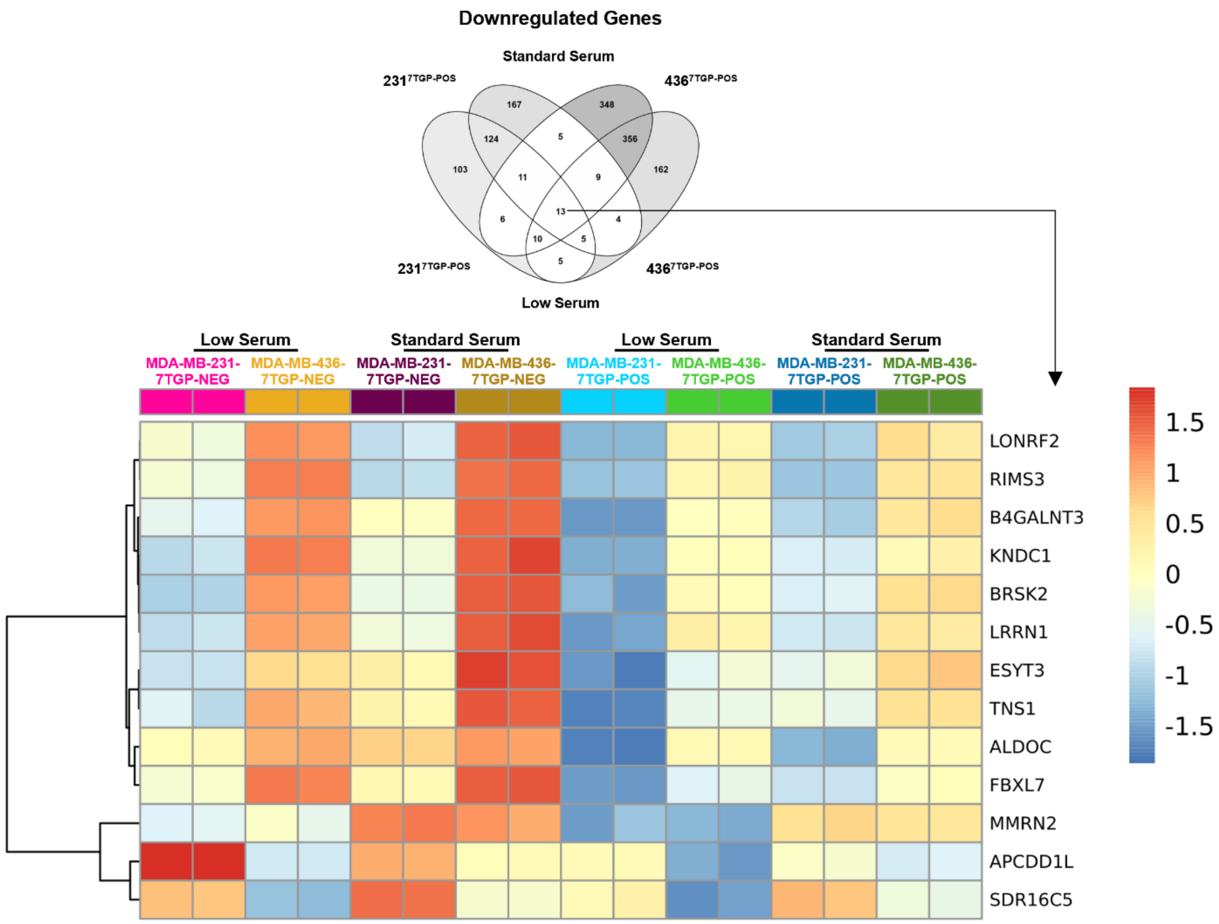


**Supplementary Fig. 7 | Shared downregulated genes across Wnt-positive TNBC cell lines and serum conditions.** Venn diagram showing the overlap of genes downregulated in Wnt-positive compared to Wnt-negative TNBC cells under both low and standard serum conditions. The heat map displays the expression patterns of the 13 genes consistently downregulated across all four comparisons. Differential expression was defined using an absolute $\log_2$ fold change ≥ 1

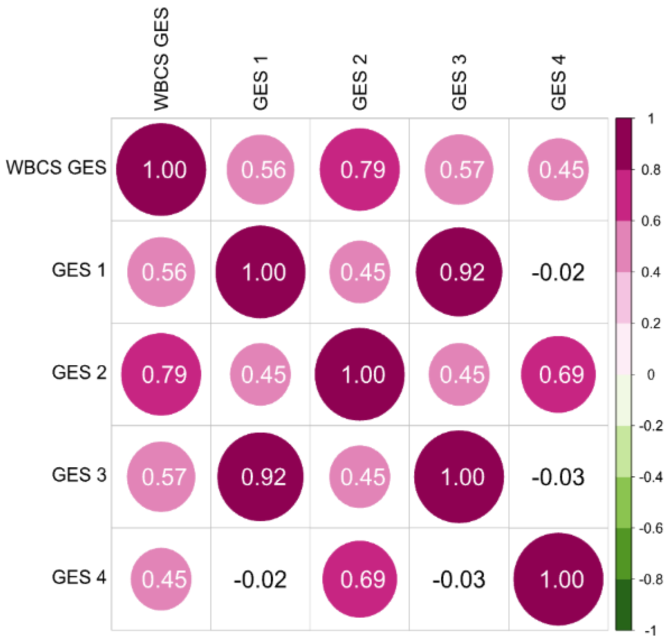


WBCS: Wnt β-catenin signature from Lehmann et al.
GES1: 687 DEG - MDA-MB-231$^{7TGP\text{-}NEG}$ vs. MDA-MB-231$^{7TGP\text{-}POS}$ (low serum)
GES2: 1,278 DEG - MDA-MB-436$^{7TGP\text{-}NEG}$ vs. MDA-MB-436$^{7TGP\text{-}POS}$ (low serum)
GES3: 848 DEG - MDA-MB-231$^{7TGP\text{-}NEG}$ vs. MDA-MB-231$^{7TGP\text{-}POS}$ (standard serum)
GES4: 1,418 DEG - MDA-MB-436$^{7TGP\text{-}NEG}$ vs. MDA-MB-436$^{7TGP\text{-}POS}$ (standard serum)

**Supplementary Fig. 8 | Correlation of Wnt-related gene expression signatures.** Spearman's correlation analysis was performed to assess the relationship between the Wnt/β-catenin signature (WBCS) and four Wnt-related gene expression signatures (GES1–GES4). The matrix displays Pearson correlation coefficients, indicating the strength of association between signatures.

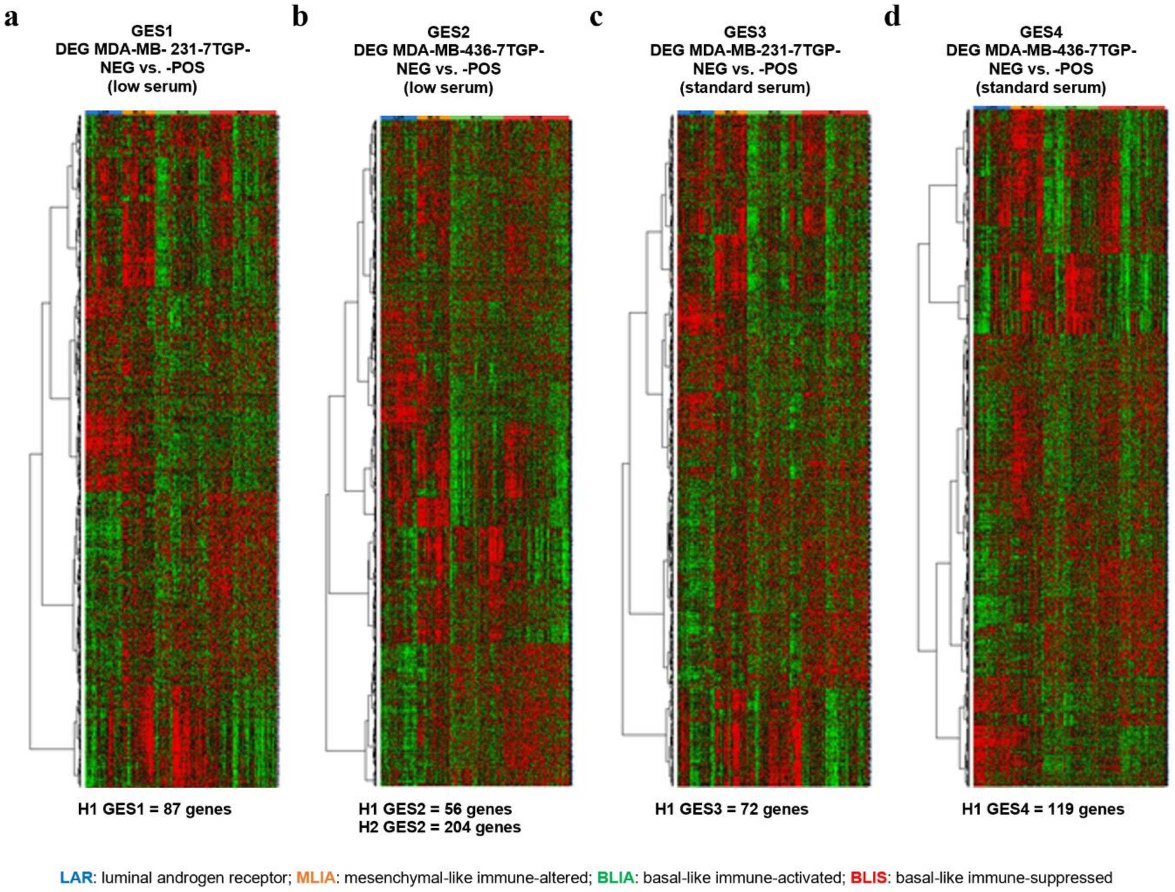


**Supplementary Fig. 9 | Expression of Wnt-related gene signatures across TNBC molecular subtypes.** Heat maps showing the expression of four Wnt-related gene expression signatures (GES1–GES4), derived from comparisons between Wnt-positive (POS) and Wnt-negative (NEG) TNBC cell lines, in a TNBC RNA-Seq cohort (n = 699). Samples are categorized into four molecular subtypes: LAR (n = 137), MLIA (n = 121), BLIA (n = 199), and BLIS (n = 242). **a**, GES1: 687 DEGs from MDA-MB-231$^{7TGP\text{-}POS}$ vs. MDA-MB-231$^{7TGP\text{-}NEG}$ under low serum. **b**, GES2: 1,278 DEGs from MDA-MB-436$^{7TGP\text{-}POS}$ vs. MDA-MB-436$^{7TGP\text{-}NEG}$ under low serum. **c**, GES3: 848 DEGs from MDA-MB-231$^{7TGP\text{-}POS}$ vs. MDA-MB-231$^{7TGP\text{-}NEG}$ under standard serum. **d**, GES4: 1,418 DEGs from MDA-MB-436$^{7TGP\text{-}POS}$ vs. MDA-MB-436$^{7TGP\text{-}NEG}$ under standard serum. Gene expression was clustered using hierarchical clustering with centered Pearson's correlation and average linkage. The number of genes in each cluster is indicated below each panel.

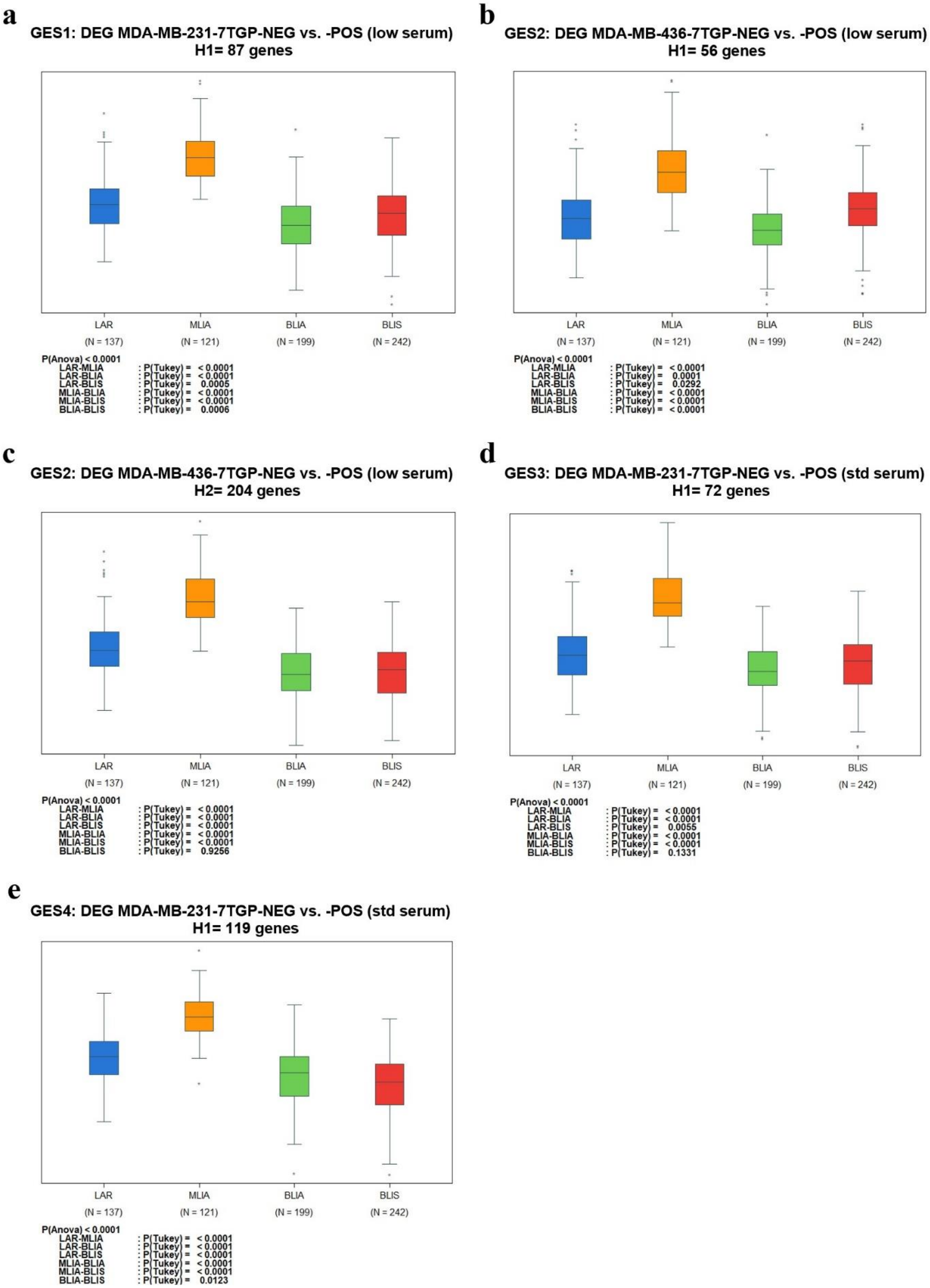
a
GES1: DEG MDA-MB-231-7TGP-NEG vs. -POS (low serum)
H1= 87 genes
LAR (N = 137)
MLIA (N = 121)
BLIA (N = 199)
BLIS (N = 242)
P(Anova) < 0.0001
LAR-MLIA : P(Tukey) = < 0.0001
LAR-BLIA : P(Tukey) = < 0.0001
LAR-BLIS : P(Tukey) = 0.0005
MLIA-BLIA : P(Tukey) = < 0.0001
MLIA-BLIS : P(Tukey) = < 0.0001
BLIA-BLIS : P(Tukey) = 0.0006
b
GES2: DEG MDA-MB-436-7TGP-NEG vs. -POS (low serum)
H1= 56 genes
LAR (N = 137)
MLIA (N = 121)
BLIA (N = 199)
BLIS (N = 242)
P(Anova) < 0.0001
LAR-MLIA : P(Tukey) = < 0.0001
LAR-BLIA : P(Tukey) = 0.0001
LAR-BLIS : P(Tukey) = 0.0292
MLIA-BLIA : P(Tukey) = < 0.0001
MLIA-BLIS : P(Tukey) = < 0.0001
BLIA-BLIS : P(Tukey) = < 0.0001
c
GES2: DEG MDA-MB-436-7TGP-NEG vs. -POS (low serum)
H2= 204 genes
LAR (N = 137)
MLIA (N = 121)
BLIA (N = 199)
BLIS (N = 242)
P(Anova) < 0.0001
LAR-MLIA : P(Tukey) = < 0.0001
LAR-BLIA : P(Tukey) = < 0.0001
LAR-BLIS : P(Tukey) = < 0.0001
MLIA-BLIA : P(Tukey) = < 0.0001
MLIA-BLIS : P(Tukey) = < 0.0001
BLIA-BLIS : P(Tukey) = 0.9256
d
GES3: DEG MDA-MB-231-7TGP-NEG vs. -POS (std serum)
H1= 72 genes
LAR (N = 137)
MLIA (N = 121)
BLIA (N = 199)
BLIS (N = 242)
P(Anova) < 0.0001
LAR-MLIA : P(Tukey) = < 0.0001
LAR-BLIA : P(Tukey) = < 0.0001
LAR-BLIS : P(Tukey) = 0.0055
MLIA-BLIA : P(Tukey) = < 0.0001
MLIA-BLIS : P(Tukey) = < 0.0001
BLIA-BLIS : P(Tukey) = 0.1331
e
GES4: DEG MDA-MB-231-7TGP-NEG vs. -POS (std serum)
H1= 119 genes
LAR (N = 137)
MLIA (N = 121)
BLIA (N = 199)
BLIS (N = 242)
P(Anova) < 0.0001
LAR-MLIA : P(Tukey) = < 0.0001
LAR-BLIA : P(Tukey) = < 0.0001
LAR-BLIS : P(Tukey) = < 0.0001
MLIA-BLIA : P(Tukey) = < 0.0001
MLIA-BLIS : P(Tukey) = < 0.0001
BLIA-BLIS : P(Tukey) = 0.0123

**Supplementary Fig. 10 | H1 and H2 clusters from Wnt-related gene signatures are differentially enriched across TNBC subtypes.** H1 and H2 gene clusters derived from Wnt-related gene expression signatures were evaluated in a TNBC RNA-Seq cohort (n = 699) comprising four molecular subtypes: luminal androgen receptor (LAR), mesenchymal-like immune-altered (MLIA), basal-like immune-activated (BLIA), and basal-like immune-suppressed (BLIS). Clusters were identified by hierarchical clustering using centered Pearson's correlation. Bar plots show enrichment scores for each cluster across TNBC subtypes: **a**, H1 GES1 (87 genes), from MDA-MB-231$^{7TGP\text{-}POS}$ vs. MDA-MB-231$^{7TGP\text{-}NEG}$ under low serum. **b**, H1 GES2 (56 genes), and **c**, H2 GES2 (204 genes), from MDA-MB-436$^{7TGP\text{-}POS}$ vs. MDA-MB-436$^{7TGP\text{-}NEG}$ under low serum. **d**, H1 GES3 (72 genes), from MDA-MB-231$^{7TGP\text{-}POS}$ vs. MDA-MB-231$^{7TGP\text{-}NEG}$ under standard serum. **e**, H1 GES4 (119 genes), from MDA-MB-436$^{7TGP\text{-}POS}$ vs. MDA-MB-436$^{7TGP\text{-}NEG}$ under standard serum. Statistical significance was assessed using one-way ANOVA with Tukey's post hoc test. P values ≤ 0.01 were considered significant and are shown below each panel.

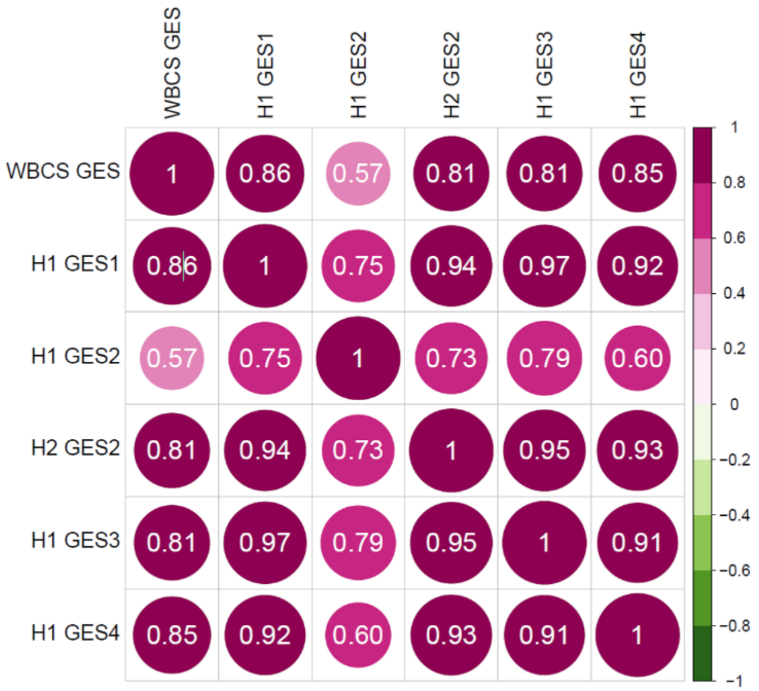


WBCS: Wnt β-catenin signature from Lehmann et al.
H1 GES1: 82 genes - MDA-MB-231$^{7TGP\text{-}NEG}$ vs. MDA-MB-231$^{7TGP\text{-}POS}$ (low serum)
H1 GES2: 56 genes- MDA-MB-436$^{7TGP\text{-}NEG}$ vs. MDA-MB-436$^{7TGP\text{-}POS}$ (low serum)
H2 GES2: 204 genes - MDA-MB-436$^{7TGP\text{-}NEG}$ vs. MDA-MB-436$^{7TGP\text{-}PO}$S (low serum)
H1 GES3: 72 genes - MDA-MB-231$^{7TGP\text{-}NEG}$ vs. MDA-MB-231$^{7TGP\text{-}POS}$ (standard serum)
H1 GES4: 119 genes - MDA-MB-231$^{7TGP\text{-}NEG}$ vs. MDA-MB-231$^{7TGP\text{-}POS}$ (standard serum)

**Supplementary Fig. 11 | Correlation of H1 and H2 clusters from Wnt-driven gene signatures with the Wnt/β-catenin signature.** Spearman's correlation analysis was used to assess the relationship between the Wnt/β-catenin signature (WBCS) and five Wnt-related H1 and H2 gene clusters derived from hierarchical clustering of gene expression in a TNBC RNA-Seq cohort (n = 699). The matrix displays Pearson correlation coefficients, reflecting the degree of association between each cluster and the WBCS.
